# Epitope-anchored contrastive transfer learning for paired CD8^+^ cell receptor-antigen recognition

**DOI:** 10.1101/2024.04.05.588255

**Authors:** Yumeng Zhang, Zhikang Wang, Yunzhe Jiang, Dene R. Littler, Mark Gerstein, Anthony W. Purcell, Jamie Rossjohn, Hong-Yu Ou, Jiangning Song

## Abstract

Understanding the mechanisms of T-cell antigen recognition that underpin adaptive immune responses is critical for the development of vaccines, immunotherapies, and treatments against autoimmune diseases. Despite extensive research efforts, the accurate identification of T cell receptor (TCR)-antigen binding pairs remains a significant challenge due to the vast diversity and cross-reactivity of TCRs. Here, we propose a deep-learning framework termed Epitope-anchored Contrastive Transfer Learning (EPACT) tailored to paired human CD8^+^ TCRs from single-cell sequencing data. Harnessing the pre-trained representations and the contrastive co-embedding space, EPACT demonstrates state-of-the-art model generalizability in predicting TCR binding specificity for unseen epitopes and distinct TCR repertoires, offering potential values for practical outcomes in real-world scenarios. The contrastive learning paradigm achieves highly precise predictions for immunodominant epitopes and facilitates interpretable analysis of epitope-specific T cells. The TCR binding strength predicted by EPACT aligns well with the surge in spike-specific immune responses targeting SARS-CoV-2 epitopes after vaccination. We further fine-tune EPACT on TCR-epitope structural data to decipher the residue-level interactions involved in T-cell antigen recognition. EPACT not only exhibits superior capabilities in quantifying inter-chain distance matrices and identifying contact residue pairs but also corroborates the presence of molecular mimicry across multiple tumor-associated antigens. Together, EPACT can serve as a useful AI approach with significant potential in practical applications and contribute toward the development of TCR-based diagnostics and immunotherapies.

## Introduction

CD8^+^ T cells play a central role in the immune system against viral infections, autoimmune diseases, and cancers that differentiated cytotoxic T lymphocytes (CTLs) can kill target cells^1–5^. TCRs composed of multiple protein chains can trigger the activation of CD8^+^ T cells by recognizing antigens presented by major histocompatibility complex (MHC) class I molecules^6, 7^. The accurate and high-throughput identification of TCR sequences that bind to specific antigens is increasingly critical for explorations of the mechanisms of T cell immune responses and to underpin the development of effective TCR-based immunotherapies^8^. In addition, binding specificities of TCR repertoires can provide an alternative to cancer diagnostic markers^9^ and to monitor the effectiveness of tumor treatment or vaccines^10, 11^.

Recent advances in single-cell sequencing techniques enable the pairing of TCRα and TCRβ transcripts through fluorescence-activated cell sorting (FACS) isolation or emulsion-based methods^12^. Despite the lower throughput than bulk TCR sequencing methods, capturing paired TCRαβ information is bound to promote the characterization of TCR diversity and function. Various experimental approaches, such as tetramer-associated TCR sequencing^13^ (TetTCR-seq) and microfluidic antigen-TCR engagement sequencing^14^ (MATE-seq), were developed for the mapping of paired TCRαβ sequences to antigen recognition specificity at the single-cell level. However, these powerful experimental methods have several shortcomings, including high cost, technical complexity, and limited epitope coverage^12^. On the other hand, TCR cross-reactivity^15^ that one TCR can bind to multiple peptide-MHC (pMHC) complexes presents therapeutic potentials to devise T cells with cross-reactive TCRs targeting various tumor antigens^16^ yet can provoke risky autoimmune responses when T cells respond to self-antigens^17^. Molecular mimicry between activated peptides and the plasticity of complementarity-determining regions (CDRs) can jointly contribute to TCR cross-reactivity^18, 19^. Still, the availability of the TCR-pMHC complex crystal structures is far from underpinning systematic investigations of the intricate mechanism. Addressing current challenges in TCR-antigen recognition^20^, it is necessary to exploit state-of-the-art AI systems to predict binding specificity and interaction conformation between TCR and pMHC complex.

A multitude of computational approaches pinpoint a promising direction to tackle the issue of TCR-antigen binding specificity via cutting-edge deep learning frameworks^21^. Existing methods comprise three major categories: (1) TCR representation models (GLIPH2^22^, DeepTCR^23^, TCRdist3^24^, TCR-BERT^25^); (2) peptide-specific TCR binding models (TCRex^26^, TCRGP^27^, NetTCR-2.0^28^, TCRAI^29^, MixTCRpred^30^); (3) pan-specific TCR binding models (ERGO-II^31^, TITAN^32^, pMTnet^33^, TEIM-Seq^34^, PanPep^35^, STAPLER^36^, TAPIR^37^, TULIP-TCR^38^, NetTCR-2.2^39^, pMTnet-omni^40^), but most of these works only consider the CDR3 loop of the TCRβ chain. Despite the dominant role of CDR3β in antigen recognition and TCR diversity, the TCRα chains also contact the pMHC complexes and contribute to the interaction, such that pairing inputs of TCRαβ sequences should provide a more comprehensive view of TCR binding specificity^41^. Besides, pan-specific models that embed TCR and pMHC sequences simultaneously are designed to generalize to neoantigens or other less common peptides. However, few analyses include evaluation under zero-shot settings^35^, resulting in the over-optimistic performance of state-of-the-art predictors. Model capacities, especially those handling paired TCRαβ sequences, are still far from satisfactory. Moreover, the lack of high-quality negative data and false-negative pairs from biased data generation also hinders AI applications in real-world scenarios^42^. To decipher the underlying binding mechanisms from a structural perspective, TEIM-Res first harnessed deep learning techniques to predict the pairwise residue interactions between CDR3β and epitope sequence^34^. Nevertheless, other CDR loops, such as the CDR1 and CDR3 of the TCR alpha chain, are also involved in the structural interplay between TCR and epitope^43^, and no existing computational methods concern the in-depth analysis of TCR cross-reactivity.

Here, we propose a deep-learning framework, epitope-anchored contrastive transfer learning (EPACT), for paired CD8^+^ T cell receptor-antigen recognition. Leveraging the contextualized representations from the pre-trained language model^25, 36^ and the prior pMHC binding/presentation embeddings^33, 40^, EPACT achieves robust adaptivity to novel TCR-pMHC pairs through transfer learning. Meanwhile, supervised contrastive learning adopting epitope/pMHC anchors preserves the prediction specificity for a particular epitope and provides an interpretable co-embedding space for TCRs and cognate pMHC targets^44^. We evaluate the model generalizability under two scenarios for binding specificity prediction: (1) predicting binding TCR for unseen epitopes and (2) adapting to distinct TCR populations. In addition to state-of-the-art performance in distinguishing pairing TCRs of given pMHC complex from decoys, EPACT also exhibits outstanding capacity in illuminating the residue-level interactions within the CDR-epitope interface. We further apply EPACT to SARS-CoV-2 epitope-specific TCR clonotypes under diverse infection and vaccination conditions^45^ as well as structure-driven TCR cross-reactivity instances in autoimmune diseases^46^ and cancer immunotherapies^47^. Our analyses demonstrate the application potential of EPACT in accelerating the development of TCR-based diagnostics and immunotherapies for infectious diseases and cancers.

## Results

### Overview of the EPACT methodology

We employed a divide-and-conquer paradigm to develop the architecture of EPACT, concentrating on the interaction between paired TCRαβ chains from CD8^+^ T cells and the cognate peptide-MHC targets (**Fig. 1a,b**). The detailed model architecture is presented in **Supplementary Fig. 1**. Specifically, we first pre-trained separate protein language models^48^ that reconstructed masked amino acids and Atchley factors^49^ for TCR or peptide sequences. Transformer-based language models harnessed the vast collection of immune epitopes and diverse TCR repertoires, thus yielding contextualized embeddings for CD8^+^ T cell epitopes and receptors. We employed residual convolutional blocks^50^ to encode the evolutionary and biophysical properties of MHC alleles, as MHC class I molecules present the epitopes to TCR on the cell surface^51^. We then combined the MHC features with prior peptide embeddings to learn the fused representation of the pMHC complex via a dual cross-attention (DCA) module. After incorporating MHC information, a language modeling head was inherited from the peptide language model to predict masked amino acids along the peptide sequences. We trained a peptide-MHC binding model on binding affinity data collected from NetMHCpan-4.1^52^. The predicted normalized IC_50_ values of test pMHC pairs were highly correlated with the experimental measures across multiple HLA gene subtypes (**Fig. 1e** and **Extended Data Fig. 1a**), with an overall Pearson correlation coefficient of 0.822. We also assessed an epitope presentation model using the independent test set of BigMHC^53^. Our intermediate model significantly improved the prediction of MHC class I eluted ligands (**Fig. 1f** and **Extended Data** Fig. 1b-d), achieving a mean area under the precision-recall curve (AUPR) of 0.901 when stratifying by MHC alleles (BigMHC AUPR: 0.878, NetMHCpan-4.1 AUPR: 0.831).

**Fig. 1.**
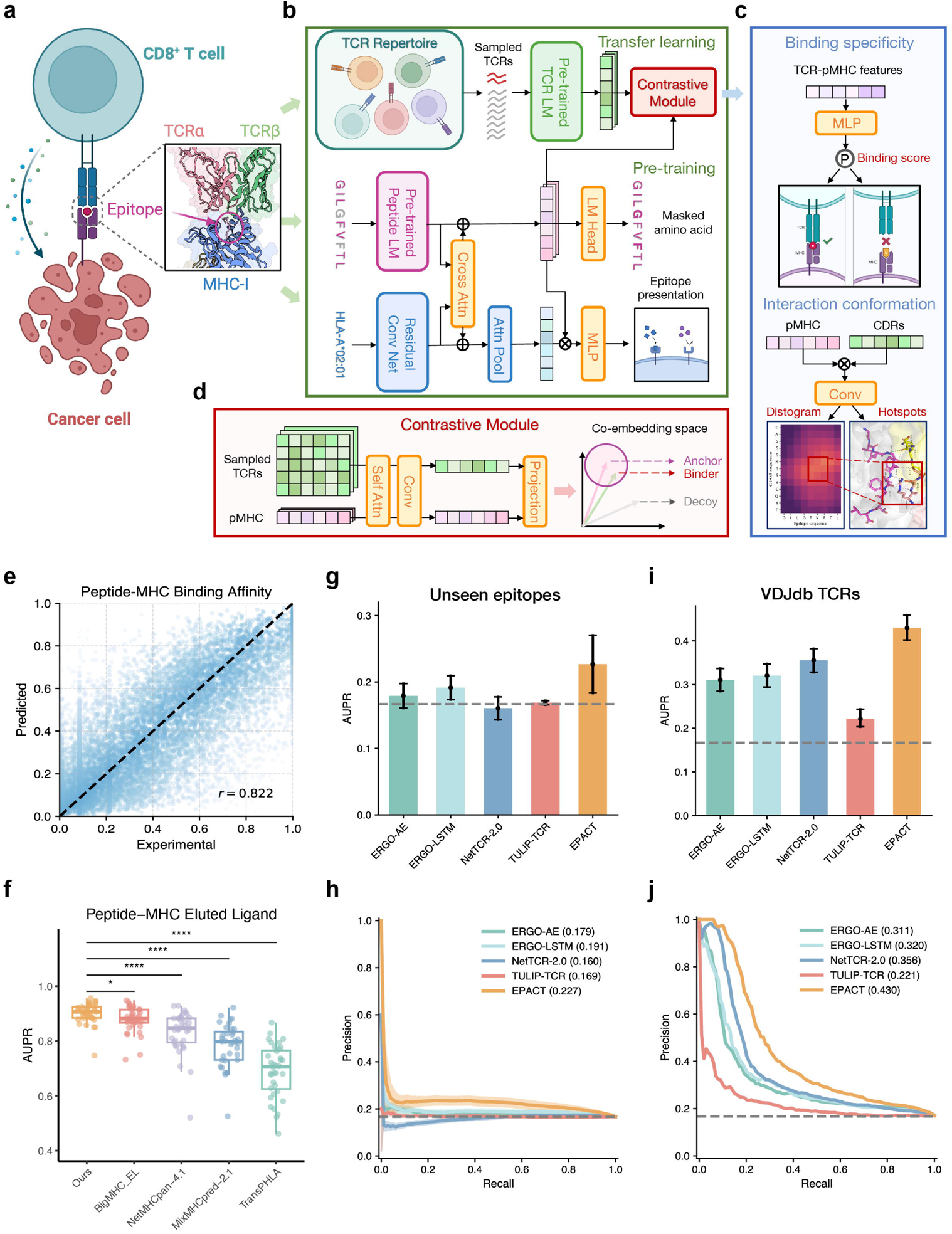
EPACT boosts CD8+ TCR-pMHC recognition for unseen peptides and distinct TCR populations. **a,** Schematic diagram of TCR-pMHC recognition. Created with BioRender.com. **b,** The model backbone of EPACT consists of pre-trained peptide and TCR language models, a pMHC model to predict epitope binding or presentation, and a contrastive learning module. The epitope representations with fused MHC information and the sampled TCR embeddings are fed into the contrastive learning module together. **c,** Two related tasks for predicting TCR-pMHC recognition: (top) Binding specificity prediction, output a binding score to decide whether the input TCR-pMHC pairs can bind together; (bottom) Interaction conformation prediction, output residue-level distance matrix and contact matrix between CDR loops and the epitope. **d,** Contrastive learning module, TCR and pMHC classification embeddings are projected into a shared latent space after feature extraction by paralleled self-attention layer and residual convolutional blocks. The contrastive loss is computed according to the cosine distances between the pMHC anchor and binding or decoy TCRs in the co-embedding space. **e,** The experimental and predicted binding affinity (normalized IC_50_ values) of test peptide-MHC pairs. **f,** Predicted AUPRs across 36 MHC alleles evaluated on the test dataset of BigMHC. The P-values were calculated by the Wilcoxon signed rank test. *P<0.05, ****P<0.0001. **g,** Bar plots of AUPRs and **h,** precision-recall curves of the candidate methods in cross-validation (predicting for unseen epitopes). The error bars indicate the standard deviation of AUPRs across five-folds, and the shade around the precision-recall curves represents the standard error of precisions. **i,** Bar plots of AUPRs and **j,** precision-recall curves of the candidate methods in testing (predicting for VDJdb TCRs). The error bars indicate the 95% confidence intervals by 1000 bootstrap iterations. The gray dashed lines denote the results of random predictions.

Leveraging the robust representations derived from powerful TCR and pMHC pre-trained models, EPACT generalized to predict TCR antigen recognition tasks via transfer learning (**Fig. 1c**). We prepared a pool of epitope-specific TCRs in the training data and then devised a contrastive learning module to connect the TCR and pMHC subnetworks (**Fig. 1d**): 1) For each TCR-pMHC pair with known binding specificity, “non-binding” TCRs were randomly sampled from the TCR pool; 2) TCR and pMHC pre-trained embeddings were processed by paralleled self-attention layers and convolutional blocks with dropout; 3) Classification embeddings representing the CLS token of TCR and pMHC were subsequently projected into a co-embedding space; 4) A supervised contrastive loss^54^ was calculated to shorten the cosine distance between the pMHC anchor and binding TCR compared with non-binding ones. The classification embeddings were also concatenated to output a pan-epitope binding score ranging from 0 to 1 by a multi-layer perceptron (MLP). In addition to predicting TCR-pMHC binding specificity, we also fine-tuned EPACT to characterize the residue-residue interactions between CDR loops and the epitope. The outer product of the residue-level embeddings of TCR and epitope sequences was further fed in a two-dimensional convolutional layer to simultaneously predict distance matrix and contact residue pairs. Therefore, EPACT could identify the binding hotpots, indicating the interaction conformation of the TCR-pMHC complex.

### EPACT achieves state-of-the-art performance for predicting TCR-pMHC binding specificity

We adopted two evaluation settings to mimic the real-world applications of the TCR-pMHC binding specificity model. We clustered the training epitopes according to the pairwise sequence similarity and allocated the corresponding TCR-pMHC pairs of each cluster in different training folds. In this way, we conducted a five-fold cross-validation to assess the zero-shot predictions in which the model was expected to adjust the binding preferences of unseen peptides. It could partly reflect the model’s capacity to identify neoantigen-specific TCR clones. Genetic rearrangement during T cell development results in diverse TCR populations of different individuals, while antigen exposure and environmental factors also contribute to variations in T cell repertoires^55^. We selected the unique TCR-pMHC pairs in the VDJdb database^56^ to build the test dataset and generated mismatched TCRs in the subset. We took the average of the predicted binding scores derived from five training folds for benchmarking. Despite the confounding factors of experimental techniques and collection criteria, it was an effective way to examine model generalizability to distinct TCR populations.

The hypervariable CDR3 loops play a crucial role in antigen recognition^7^, so we only considered CDR3αβ sequences from the paired TCR chains at first. EPACT substantially enhanced model performance on unseen epitopes with paired CDR3αβ and pMHC inputs compared to other deep-learning methods (**Fig. 1g,h**). While other methods (ERGO-II^31^, NetTCR-2.0^28^, and TULIP-TCR^38^) struggled with surpassing random predictions, EPACT obtained an average AUC of 0.609 and AUPR of 0.227 across five folds. We also compared the AUCs and AUPRs when stratifying by epitopes in cross-validation. EPACT demonstrated state-of-the-art performance for the majority of epitopes (**Supplementary** Fig. 2) in which the zero-shot AUCs and AUPRs excelled those derived from NetTCR-2.0 predictions for 85.0% and 80.4% of the epitope targets. We then assessed the model generalizability on VDJdb unique TCR-pMHC pairs (**Fig. 1i,j**). All candidate models outperformed unseen epitope predictions except TULIP-TCR, an unsupervised method that transferred little knowledge to a distinct TCR population. EPACT reached a median AUC of 0.699 [95% confidence interval (CI), 0.682 to 0.717] and a median AUPR of 0.430 (CI, 0.402 to 0.459) by 1000 bootstrap iterations, while the second best method in our experiment, NetTCR-2.0, obtained a median AUC of 0.643 (CI, 0.624 to 0.661) and a median AUPR of 0.356 (CI, 0.328 to 0.382).

The CDR1 and CDR2 loops encoded by human TRAV/TRBV genes mainly contact the surface of HLA molecules^57^. Nevertheless, incorporating CDR1 and CDR2 sequences of TCRαβ chains enables the enhanced prediction performance for TCR binding specificity due to additional co-evolutionary information^58^. Furthermore, structural evidence revealed that CDR1α sequences in many TCR-pMHC complexes participate in the interactions with part of the peptide residues^43^, while CDR3β might even shift to avoid interactions with the peptide antigen under unconventional docking^59^. Therefore, we determined to extract both CDR1 and CDR2 loops from IMGT annotated V-genes^60^ and integrated them into the large language-based TCR model. Several existing methods also provided models accommodating the inputs of all six CDR loops (NetTCR-2.2^39^ and MixTCRpred^30^), CDR3 sequences plus categorical V-and J-genes (ERGO-II^31^), or full-length TCRαβ sequences (STAPLER^36^).

We conducted cross-validation and independent testing via the same setting on TCR-pMHC pairs with paired CDR3αβ sequences and annotated V-, J-genes. We also trained a model solely on CDR3αβ to assess possible improvements brought by adding CDR1 and CDR2 sequences. The zero-shot validation performance on unseen peptides showed a minimal difference between the CDR3αβ and TCRαβ model (average AUC, 0.597 vs. 0.595; average AUPR, 0.218 vs. 0.224, **Fig. 2a**). In contrast, the shared V-genes across diverse TCR populations might contribute to performance improvements of the TCRαβ model (**Fig. 2a,b**) as the sequence diversity in germline-encoded CDR1 and CDR2 loops is much lower than CDR3. The median AUC by 1000 bootstrap iterations increased from 0.665 (CI, 0.647 to 0.682) to 0.697 (CI, 0.678 to 0.714), and the median AUPR rose from 0.381 (CI, 0.354 to 0.406) to 0.444 (CI, 0.416 to 0.471). EPACT also outperformed external methods, including ERGO-II, NetTCR-2.2, and STAPLER (**Extended Data Fig. 2a-d**). The zero-shot AUCs and AUPRs of EPACT transcended those derived from NetTCR-2.2 predictions for 82.1% and 74.9% of the epitope targets (**Supplementary Fig. 2**). Although the overall prediction metrics of NetTCR-2.2 approached EPACT (median AUPR, 0.425 vs. 0.444), EPACT exhibited a higher average AUPR (0.502 vs. 0.461) by stratifying the test epitopes. We analyzed the AUPRs for the epitopes with over ten binding TCRs in the test dataset. EPACT was the best predictor for 7 in 24 epitope targets (**Fig. 2c**), including the Melan-A epitope EAAGIGILTV (AUPR, 0.948), Influenza M peptide GILGFVFTL (AUPR, 0.918), and SARS-CoV-2 nucleocapsid-derived peptide SPRWYFYYL (AUPR, 0.623). To validate the model robustness on various epitope targets, we categorized the epitopes in the validation set into groups of different sequence lengths and Levenshtein distances to training ones. Consequently, we observed EPACT’s reliability and superiority across all categories (**Supplementary Fig. 3**). We also conducted an ablation study in regards to EPACT’s essential modules, including the pre-training process and contrastive loss (**Supplementary Table 1**). The presented benchmarking results illustrate that EPACT exhibited an excellent capability of predicting CD8^+^ TCR-pMHC recognition for unseen peptides and distinct TCR populations.

**Fig. 2.**
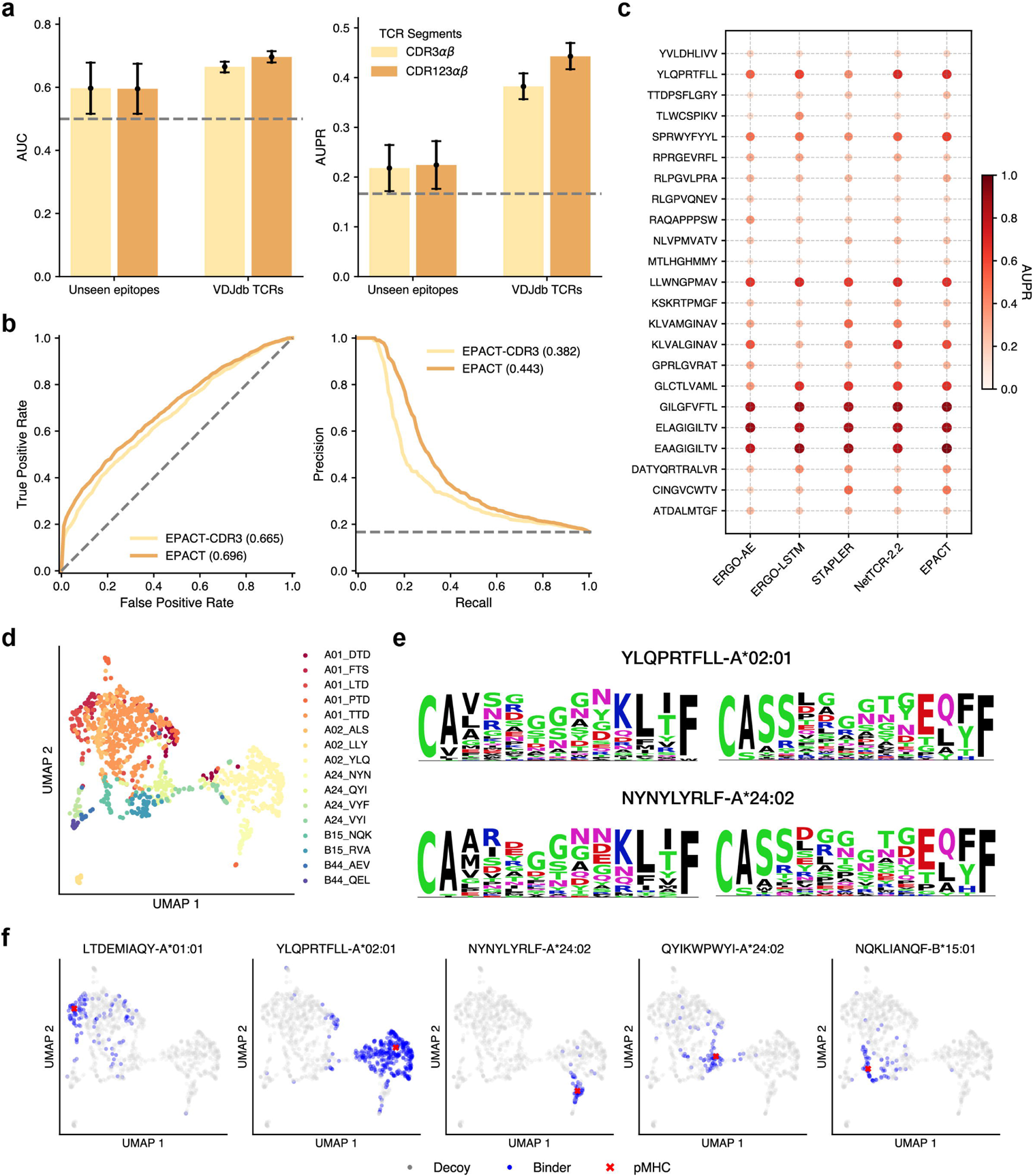
EPACT incorporates all CDR loops and interprets TCR specificity in co-embedding space. **a,** Bar plots showing EPACT’s performance in predicting binding specificity for unseen epitopes and VDJdb TCRs using different datasets that used CDR3αβ or all six CDR loops to represent TCR sequences. (left) AUC (right) AUPR. The error bars under the evaluation settings of unseen epitopes denote the standard deviation across five folds. **b,** Receiver-operating characteristic curve (left) and precision-recall curve (right) to evaluate the testing performance of EPACT-CDR3 and EPACT on VDJdb TCR-pMHC pairs. The gray dashed lines denote the results of random predictions. **c,** Comparison of AUPRs derived from ERGO-II, STAPLER, NetTCR-2.2, and EPACT for 24 epitopes with over ten binding TCRs in the test dataset. The darker color and larger size of the point indicate a higher AUPR. **d,** UMAP projection of the predicted SARS-CoV-2 epitope-specific TCR clusters. The TCR embeddings were derived from the co-embedding space via contrastive learning. **e,** Sequence motifs of CDR3α and CDR3β representing the epitope-specific TCRs for two spike protein epitopes (top) YLQPRTFLL and (bottom) NYNYLYRLF. **f,** UMAP projections of five spike epitope targets and experimental binding TCRs. The red cross in each subplot denotes the corresponding pMHC anchor, the blue points represent the binding TCRs, while the gray points indicate the decoys TCRs with no experimental evidence responding to the target, respectively.

### EPACT enables interpretable analysis of epitope-specific TCRs

Accurate identification of TCRs of particular tumor-associated or viral epitopes can help expedite vaccine development and T cell-based immunotherapies^61–63^. Previous unsupervised clustering methods, such as GLIPH2^22^ and TCRdist3^24^, mapped the input single or paired TCRs to unique clusters based on the sequence features. TCR specificities were assigned based on the resemblance to TCRs with known binding targets. However, these epitope-specific TCRs recognizing common pMHC complexes might not share high sequence similarity, especially the hypervariable CDR3 loops, partly due to the inherent diversity of TCR repertoires and TCR degeneracy^64^ (i.e., one TCR can bind to multiple antigen peptides). These present challenges for inferring the epitope-specific TCR clones among a TCR repertoire.

Contrastive learning module in EPACT mapped pMHC anchors and TCRs into an interpretable co-embedding space. Thus, we defined epitope-specific TCRs based on their cosine distances to the corresponding pMHC anchors. We re-trained an EPACT model using the complete TCRαβ-pMHC recognition dataset and derived the projections of all available pMHCs. We assumed that epitope-specific TCRs would be organized into clusters around the centroid representing the epitope targets. To illustrate the effectiveness of our EPACT model for predicting epitope-specific TCRs in the co-embedding space, we chose 16 SARS-CoV-2 epitopes and antigen-presented MHC alleles for exemplification. We constructed the SARS-CoV-2 epitope-specific TCR clusters using an interpretable approach. We assigned the candidate TCRs to the nearest pMHC anchors and set the maximum cosine distance from the anchor to 0.4 for high specificity in each cluster. We employed the UMAP algorithm to reduce the dimension of TCR projections^65^, resulting in distinct TCR clusters in the two-dimensional space (**Fig. 2d**). Interestingly, the predicted epitope-specific TCRs for different epitope targets presented by HLA-A*01:01, HLA-B*15:01, and HLA-B*44:02 were found to be in close proximity in the UMAP space. This probably implies the impact on TCR-pMHC recognition from the HLA genotypes^66^. Next, we inspected the amino acid preferences of CDR3αβ sequences responding to five 9-mer spike protein epitopes (LTDEMIAQY, YLQTRPFLL, NYNYLYRLF, QYIKWPWYI, and NQKLIKNQF) presented by various HLA molecules (**Fig. 2e** and **Supplementary Fig. 4**). The core regions of the spike-epitope-specific CDR3αβ loops that primarily contacted the epitope were highly diverse despite the conserved motifs at N-and C-terminus^67^. Nevertheless, polar amino acids, such as Glycine(G), Asparagine(N), Serine(S), and Threonine(T) were more likely to occur at the core positions of CDR3αβ sequences. To confirm the representativeness of the epitope-specific clusters, we compared the distribution of the spike and non-spike epitope targets with their known binding TCRs (**Fig. 2f** and **Supplementary Fig. 5**). As can be seen, the UMAP projections of TCRs with experimental specificity also gathered around the cognate pMHC targets and appeared to form cluster shapes similar to the predicted ones.

### EPACT aligns well with the T cell responses to SARS-CoV-2 infection and vaccination

We further investigated the application potential of EPACT to clinical cohorts through a longitudinal study^45^. Minervina et al. profiled the SARS-Cov-2 responsive CD8^+^ T cells from samples that underwent distinct antigen exposure through scRNA-seq and scTCR-seq. We collected 4,471 unique TCR clonotypes with paired CDR3αβ sequences, V-, J-gene annotations, and antigen specificity for 16 epitope targets inferred by DNA barcoded pMHC dextramer. We removed the overlapped TCRs in the training data and randomly chose non-responsive TCRs from the combined repertoire of healthy human samples. Thus, we constructed an external TCR-pMHC recognition dataset comprising 3,540 unseen SARS-CoV-2 responsive TCR clonotypes and five times non-binding TCRs for fair benchmarking.

We evaluated the model generalizability of STAPLER^36^, NetTCR-2.2^39^, EPACT, and MixTCRpred^30^ to unseen TCR clones, respectively. Model outputs of MixTCRpred were assembled from 14 peptide-specific predictors. To validate the necessity of including CDR1 and CDR2 features, we also assessed the performance of EPACT trained on CDR3αβ data and NetTCR-2.0^28^. EPACT substantially enhanced the model capacity (**Fig. 3a,b**), achieving a median AUPR of 0.509 (CI, 0.494 to 0.524) across all SARS-CoV-2 epitopes by bootstrapping. Meanwhile, the second-best method, NetTCR-2.2, achieved a median AUPR of 0.455 (CI, 0.439 to 0.472). The median AUPR of EPACT (0.425, CI, 0.410 to 0.442) significantly decreased when solely using CDR3αβ features due to the loss of underlying co-evolutionary and biophysical information from CDR1 and CDR2 loops. We also examined the model performance for each SARS-CoV-2 epitope (**Extended Data Fig. 3a**). EPACT demonstrated a nearly equivalent predictor as MixTCRpred, an ensemble of several peptide-specific models (average AUPR, 0.382 vs. 0.398). Given abundant training data, the contrastive learning paradigm empowered EPACT with high specificity^44^ so that it even outperformed MixTCRpred in predicting TCRs binding to two immunodominant SARS-CoV-2 epitopes (TTDPSFLGRY AUPR, 0.666 vs. 0.605 and YLQPRTFLL AUPR, 0.765 vs. 0.753). In addition, EPACT delivered remarkable advantages over other pan-specific models, resulting in higher AUPRs for 12 in 14 epitope targets compared to NetTCR-2.2 (**Fig. 3c**).

**Fig. 3.**
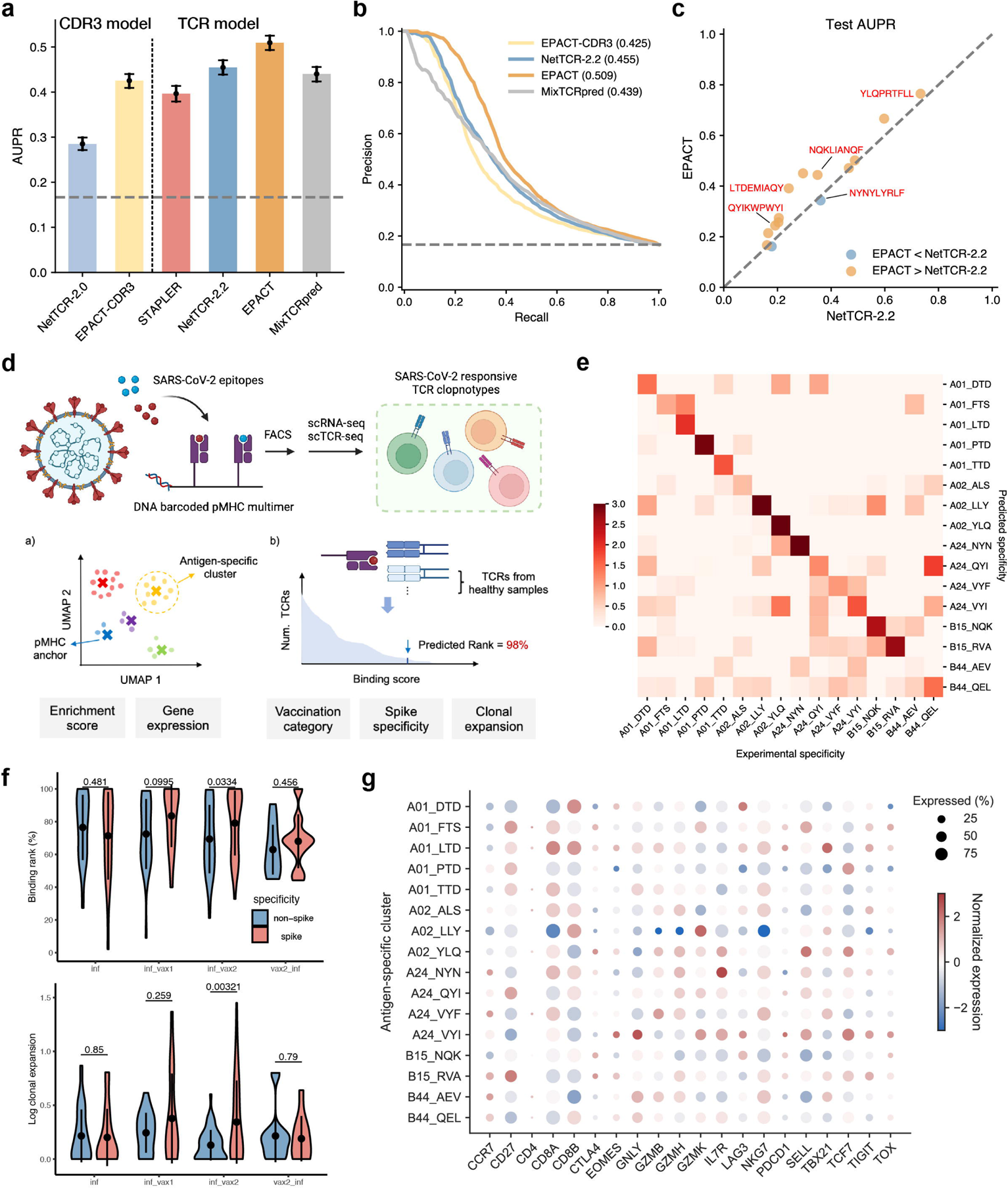
EPACT predicts epitope-specific CD8+ T-cell responses to SARS-CoV-2 infection and vaccination. **a,** Test AUPRs on unseen SARS-CoV-2 TCR-pMHC recognition dataset. Two methods receiving paired CDR3αβ inputs and another four based on additional V-, J-gene annotations were compared. **b,** Precision-recall (PR) curves of EPACT-CDR3, NetTCR-2.2, EPACT, and MixTCRpred. The gray dashed lines denote the results of random predictions. **c,** Pairwise comparison of test AUPRs for 14 epitope targets between two models using CDR1, CDR2, and CDR3 sequences (EPACT and NetTCR-2.2). Five spike epitopes are annotated in red text. **d,** Workflow to analyze SARS-CoV-2 responsive TCR clonotypes collected from Minervina et al., including antigen-specific clusters and TCR binding ranks. **e,** Heatmap of log enrichment ratios of TCRs with experimental specificity across predicted antigen-specific clusters. Darker colors along the diagonal indicate better alignment between prediction and experiment results. **f,** Median binding rank and log clonal expansion of spike-specific and non-spike-specific TCR across four groups upon diverse SARS-CoV-2 antigen exposure and vaccination (inf, inf-vax1, inf-vax2, and vax2-inf). The comparisons between spike-specific and non-spike-specific TCR responses were conducted by Student’s t-test (P-values are displayed above the violin plots). **g,** Bubble plot of normalized expressions and fractions of expressed cells of T cell mark genes in each antigen-specific TCR cluster defined by EPACT. The circle size denotes the percentage proportion of cells expressing a marker gene in each cluster, while the color scale indicates normalized gene expression across all clusters.

We next applied EPACT to the entire SARS-CoV-2 epitope-specific CD8+ T cell repertoires in the cohort to explore whether EPACT could detect the dynamics of T cell binding specificity and other phenotypes under a real-world scenario (**Fig. 3d**). We predicted antigen-specific clusters in the TCR repertoire by calculating the cosines distances between TCRs and pMHC anchors in the co-embedding space (**Extended Data Fig. 3b,c**). To determine the relative binding strength to the epitope target, we generated the background distribution of binding score by sampling a large collection of TCRs and subsequently a rank-percentile measurement. The predicted rank reflected the binding strength of query TCR relative to the background TCRs. We then monitored the variation in binding rank and TCR clonal expansion upon diverse SARS-CoV-2 antigen exposure and vaccination.

We validated the effectiveness of predicted antigen-specific TCR clusters by comparing the abundance within and outside the cluster of TCRs with experimental specificity. We calculated the enrichment ratios of various SARS-CoV-2 epitope-specific TCRs across 16 clusters (**Fig. 3e**). As a result, the antigen-specific clusters derived from EPACT predictions were consistent with experimental specificity of the majority of the epitope targets^45^. For instance, five groups of spike-epitope-specific TCRs were highly enriched in the corresponding clusters; the enrichment ratios were 7.25, 73.8, 35.3, 3.29, and 12.3 for A01_LTD, A02_YLQ, A24_NYN, A24_QYI, and B15_NQK responsive TCRs, respectively. The corresponding P-values derived from the Chi-square independent test were all smaller than 0.001. Due to the limited training data, the TCR clonotypes near the pMHC anchor representing B44_AEV or B44_QEL exhibited ambiguous experimental specificity. We also inspected the gene expression profiles of the antigen-specific TCR clusters (**Fig. 3g**), including cytotoxic markers (*NKG7*, *GNLY*, *GZMB*, *GZMH*), memory markers (*TCF7*, *IL7R*, *SELL*), and exhaustion markers (*CTLA4, PDCD1*, *TOX, TIGIT*)^68^. Although the antigen-specific T cell population was composed of T cells with various functions and phenotypes from different samples^45^, we inferred several general characteristics of the T cell composition in the antigen-specific clusters. Spike-specific T cells corresponding to A02_YLQ and A24_NYN might maintain large amounts of T cells with durable cellular memory according to the upregulated expression of the memory markers. T cells targeting A02_LLY accounted for a lower proportion of differentiated effector T cells due to the downregulated expression of cytotoxic markers, which partly resulted from reduced proportions in vaccinated samples.

We compared the binding ranks of TCR-pMHC pairs from each sample across five categories, including infection only (inf), vaccinated only (vax2), infected followed by one/two doses of vaccine (inf-vax1/inf-vax2), and breakthrough infection after two doses of vaccine (vax-inf). Median binding ranks by stratifying donors and epitopes across infection or vaccination categories revealed an increase in the binding strength with spike epitopes after vaccination compared to non-spike responses (**Fig. 3f**), especially in the inf-vax2 group (*P*=0.033, Student’s t-test). We observed a similar trend in T cell clonal expansion that the clone sizes of spike-specific TCRs were greater than non-spike clones after vaccination, especially in the inf-vax2 group (*P*=0.003, Student’s t-test), to corroborate EPACT’s predictions. We further divided the SARS-CoV-2 responsive TCR clonotypes into “Strong binder” (≥99.5%), “Weak binder” (³95%), and “Others”. Strong binders with spike epitopes and the union of strong and weak binders accounted for a higher proportion in the TCR repertoire (**Extended Data Fig. 3d-f**). Our analyses aligned well with the experimental findings that spike and non-spike T cell response varied with SARS-CoV-2 infections and mRNA vaccination.

### EPACT uncovers residue-level interactions between epitope and CDR loops

Despite the complicated recognition mechanism between paired TCR chains and the pMHC complex to trigger immune responses, the residual-level interaction undoubtedly plays an essential part in the formation and stability of the TCR-pMHC complex^7^. Accurate identification of hydrogen bonds and van der Waals interactions between epitope and CDR loops can promote understanding of the potential source of TCR degeneracy and cross-reactivity. We first cross-validated the fine-tuned version of EPACT and TEIM-Res^34^, a state-of-the-art predictor for residue-level interactions between CDR3β and epitope, on 148 public TCR-pMHC complex structures in STCRDab^69^. We also included a baseline method that output an average distance matrix. We ensembled the validation predictions of CDR3β-epitope interactions and calculated the Pearson correlation coefficient (PCC) and root mean squared error (RMSE) for distance matrix prediction and area under ROC curve (AUC) for contact site prediction (**Fig. 4a**). EPACT manifested a significant advance compared to the average baseline and TEIM-Res. Specifically, EPACT achieved a median PCC of 0.953 (TEIM-Res, 0.942, *P*=5e-7, Paired t-test), a median RMSE of 2.05 (TEIM-Res, 2.22, *P*=2e-6), and a median AUC of 0.966 (TEIM-Res, 0.958, *P*=0.05). EPACT outperformed TEIM-Res in predicting CDR3β-epitope interactions for almost 70% of the available TCR-pMHC structures (**Extended Data Fig. 4**).

**Fig. 4.**
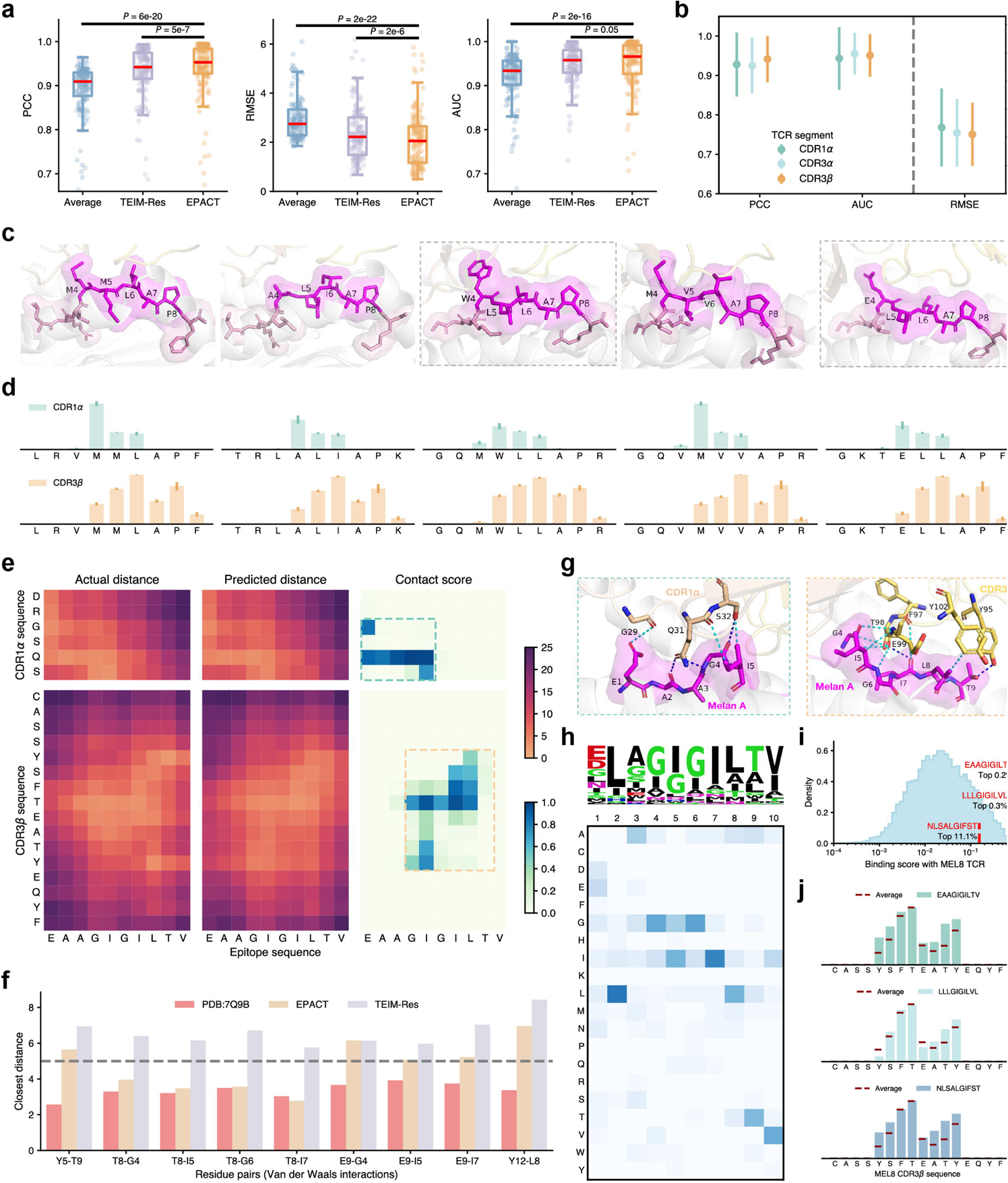
EPACT characterizes TCR-epitope interaction conformation and structure-driven TCR cross-reactivity. **a,** Box plots displaying the cross-validation PCC (left), RMSE (middle), and AUC (right) in predicting distance and contact matrix between CDR3β and epitope by average baseline, TEIM-Res, and EPACT. The P-values are derived from paired t-tests. The median values are highlighted in red. **b,** Cross-validation PCC, RMSE, AUC in predicting distance and contact matrix between CDR1α, CDR3α, or CDR3β and epitope. Higher PCC and AUC stand for better performance, whereas RMSE is the opposite. **c,** Visualization of the representative TCR-pMHC binding interfaces for YEIH^bac^, PRPF3^self^, GPER1^self^, RNASEH2B^self^, and gspD^bac^. The TCR-pMHC complex structures were retrieved from the PDB database (PDB IDs: 7N2Q, 7N2R, and 7N2P) or predicted by the TCRmodel2 sever (structures surrounded by gray dashed lines). **d,** Average contact scores to CDR1α (top) and CDR3β (bottom) loops of the amino acids along the peptide sequence predicted by EPACT. The error bars denote the standard deviations of contact scores across all activated AS or AU TCRs. **e,** Residue-residue experimental (left) and predicted (middle) distance matrices and predicted contact scores (right) characterizing CDR1α-epitope (top) and CDR3β-epitope (bottom) interactions in the MEL8 TCR-Melan A peptide-HLA-A*02:01 complex. The color scales in the heatmap represent amino acid pairs from close to distant and contact scores from low to high. The core interaction regions are surrounded by the dashed lines. **f,** Bar plots comparing the experimental distances in PDB structures (PDB ID: 7Q9B) and predicted distances by EPACT or TEIM-Res of nine inter-chain residue pairs from CDR3β and Melan A peptide. The gray dashed line indicates the contact threshold of 5 Å in our training setting. **g,** Visualization of the core interaction regions, including CDR1α (left) and CDR3β (right) loops of MEL8 TCR and Melan A peptide. The cyan and dark blue line between residues denote van der Waals interaction and hydrogen bond, respectively. **h,** Sequence motif (top) and heatmap (bottom) to display the positional amino acid preferences of peptides recognized by MEL8 TCR. **i,** Density plot to show the distribution of predicted binding scores to MEL8 TCR among the IEDB HLA-A*02:01-presented peptides. The x-axis is transformed into a log scale. **j,** Contact scores with Melan A (top), BST2 (middle), and IMP2 (bottom) peptide along the CDR3β sequence of MEL8 TCR. The red lines indicate the average contact level with the top 11.1% peptide binders along the CDR3β sequence.

Moreover, EPACT supported the investigation of interaction conformation between epitope and other CDR loops than CDR3β, thanks to incorporating all six CDR sequences. After analyzing the distances between the closest residue pairs in each TCR-pMHC complex (involvement of CDR1α and CDR3α in van der Waals interactions with the epitope in 76.4% and 86.5% of the structures), we chose to quantify the structural interplay between CDR1α, CDR3α, and the epitope (**Fig. 4b**). The cross-validation metrics for distance and contact predictions containing CDR1α and CDR3α amino acid residues were comparable to CDR3β-epitope predictions (e.g., average PCC, CDR1α: 0.928, CDR3α: 0.925, CDR3β: 0.942).

We applied the interaction model to interrogate the residue-level binding characteristics between a series of cross-reactive TCRs and their epitope targets. Yang et al. performed peptide activation assays to determine the activated human or microbial peptides presented by HLA-B*27:05 for several TCRs with a disease-associated TRBV9–CDR3β motif^46^. We predicted the distance and contact matrices for 60 activated TCR-peptide pairs to interpret TCR cross-reactivity leading to the autoimmune disease. We attempted to discover shared properties of interaction conformation at the TCR-pMHC interface. We selected the five most common peptides that activated the expanded TCR clonotypes in ankylosing spondylitis (AS) and acute anterior uveitis (AAU) patients, including YEIH^bac^ (LRVMMLAPF), PRPF3^self^ (TRLALIAPK), GPER1^self^ (GQMWLLAPR), RNASEH2B^self^ (GQVMVVAPR), and gspD^bac^ (GKTELLAPF). The structure organization of the peptides depicted a conserved TCR recognition mode emphasizing crucial contact sites at P4, P6, and P8 of the peptide^46^ (**Fig. 4c**). Contact scores predicted by EPACT not only identified the binding hotspots and structural motif (amino acid at P4 stretching out towards the CDR1α loop while amino acid side chains at P6 and P8 facing the CDR3β) but also captured the slight structural deviations (**Fig. 4d**). Methionine (M) at P4 obtained a higher contact score to the CDR1α loop than other amino acids, which coincided with structural evidence regarding side chain arrangement.

### EPACT facilitates the illumination of molecular mimicry in TCR cross-reactivity

To further explore the underlying mechanism of TCR cross-reactivity, we applied EPACT to predict the interaction conformation between a cancer-reactive MEL8 TCR and three tumor-associated epitopes in the context of HLA-A*02:01^47^. The TCR clone was derived from a stage IV malignant melanoma patient with successful tumor-infiltrating lymphocyte (TIL) therapy^70^. Dolton et al. attributed the multipronged T-cell recognition of different cancer-specific or pan-cancer antigens to molecular mimicry according to the shared binding motif and structural hotspots. Since the crystal structure of MEL8 TCR-pMHC was not included in the training data, we directly utilized EPACT to quantify the residue-residue distance matrices and predict inter-chain contact residue pairs between CDR1α, CDR3α, CDR3β loops of MEL8 TCR and a 10-mer Melan A peptide EAAGIGILTV (**Fig. 4e**). We extracted the corresponding distance maps from the complex structure (PDB: 7Q9B) to validate the predicted interactions. Amino acids of CDR3α were less likely to form stable interactions with the tumor-associated peptide (closest residue-residue distance, 4.5Å, **Supplementary Fig. 6**), so we focused on CDR1α and CDR3β in the following analyses. EPACT demonstrated an outstanding prediction performance for the core regions of the CDR1α-epitope (PCC, 0.961; RMSE, 0.782; AUC, 1.00) and CDR3β-epitope (PCC, 0.728; RMSE, 1.71; AUC, 0.852) interfaces. We chose the residue pairs that probably formed van der Waals forces or hydrogen bonds (closest distance ≤ 4.0Å) from CDR1α, CDR3β, and the peptide and compared the experimental distances and predicted distances by EPACT (**Fig. 4f**,**g** and **Supplementary Fig. 6**). We also included TEIM-Res predictions for these contact amino acid pairs. EPACT significantly reduced the prediction errors for nearly all the residue pairs compared to the state-of-the-art method TEIM-Res. Specifically, EPACT reconstructed the CDR3β binding mode (with minor prediction errors) around the central Threonine (T), connecting four consecutive peptide amino acids (G4, I5, G6, and I7)^47^ by one hydrogen bond and van der Waals interactions. The consensus on core interacting sites also suggested the peptide motif contributing to the molecular mimicry.

Leveraging EPACT’s binding specificity model, we characterized the amino acid preference among the tumor-associated peptides to activate the cross-reactive MEL8 TCR. More specifically, we curated a collection of 10-mer peptides presented by HLA-A*02:01 from IEDB^71^ and predicted their binding scores with the MEL8 TCR. We randomly chose 2,000 peptide sequences from the background population and utilized a simulated annealing strategy to generate the peptide motif (**Fig. 4h**). We introduced a point mutation to each peptide sequence and updated its binding score to the TCR target in one iteration. After filtering the favorable mutations for 500 iterations, the amino acid preference derived from the top 2% of predictions successfully captured the G-I-G-I motif, similar to positional scanning in the experimental peptide library^72, 73^. *In silico* simulation also implicitly suggests a possible position shift of the G-I-G-I motif, while the Melan A_A2L_ peptide ELAGIGILTV might initiate MEL8 T cell activation even more effectively. The Melan A peptide, bone marrow stromal antigen 2 (BST2) peptide LLLGIGILVL, and insulin-like growth factor 2 mRNA-binding protein 2 (IMP2) peptide NLSALGIFST that responded to MEL8 TCR in activation assays obtained top binding ranks (top 0.2%, 0.3%, and 11.1%) among the IEDB HLA-A*02:01 presented peptides (**Fig. 4i**). These peptides also demonstrated elevated contact levels with the side regions of CDR1α and CDR3β loops compared to the average level of other top binders (**Fig. 4j**). In addition, we employed the validation prediction for interactions between another cross-reactive TCR (MEL5) and the BST2 peptide (**Extended Data Fig. 5a-e**) to affirm EPACT’s capacity to decipher the driving factors of molecular mimicry. In addition, structural modeling results of MEL8-BST2 peptide and MEL8-IMP2 peptide interactions also provided additional evidence for the EPACT-predicted interaction conformations (**Supplementary Fig. 7**).

## Discussion

EPACT represents a novel interpretable framework to address the multi-scale interaction within the TCR-pMHC complex and adapt to emerging paired TCR sequencing data by single-cell techniques. It has achieved state-of-the-art performance in predicting TCR binding specificity and residue-level contacts, leveraging the power of a pre-trained language model and contrastive learning. We have also demonstrated the application potential of EPACT by performing in-depth analysis, including identification of antigen-specific TCR clusters in T cell repertoire, estimation of SARS-CoV-2 spike and non-spike specific T cell response upon vaccination, and investigation of TCR cross-reactivity after immunotherapy to recognize multiple tumor-associated antigens. In accordance with the sustained release of high-quality TCR binding specificity data and TCR-pMHC complex structures, EPACT is expected to be developed and explored as a more practical computational tool to accelerate disease diagnosis and assessment of T cell-based immunotherapies and vaccines in diverse clinical studies.

Despite the superiority of EPACT over other methods in model generalizability and interpretability, it still can be improved to be applied to the real-world clinical scenario, especially for predicting the responsive TCRs for neoepitopes. For predicting TCR antigen recognition for other targets with fewer binding TCRs, the commonly used negative sampling strategy might introduce biases that cannot be ignored^42^. Data scarcity of the less frequent HLA alleles might also influence the model predictions^21^. Although EPACT has already provided a version that accepts only CDR3αβ sequences of TCRs to handle the missing V-, J-gene annotations, it cannot deal with other partial inputs, such as single TCR chain and multiple TCR alpha chains that still often exist in scTCR-seq data^30^ or lacking MHC allele information. Excluding these incomplete records might affect EPACT’s performance in linking TCR binding specificity with T cell expression and phenotypes derived from diverse antigen exposure and clinical treatments. In addition to the challenges caused by data imbalance, we plan to extend the interaction conformation predicted by EPACT to reconstruct the three-dimensional structure of the CDR-epitope interface^74^ in our future work, which hopefully will provide a more intuitive view to model and interpret molecular mimicry and TCR cross-reactivity.

## Methods

### Datasets

#### Pre-training peptides and TCRαβ sequences

To prepare the pre-training dataset of human peptides, we filtered the linear epitope sequences within the length of 8-25 amino acids of the positive T cell and MHC ligand assays in the immune epitope database^71^ (IEDB). The pre-training corpus of paired TCRαβ sequences was obtained from the single-cell immune profiling data of the healthy and tumor donors in 10X Genomics Datasets (https://www.10xgenomics.com/resources/datasets, **Supplementary Table 2**) and five previous studies adopted in STAPLER^75–79^. All unique TCR clonotypes with CDR3 sequence, V-and J-gene annotations were extracted for TCR alpha and beta chain. Next, 1,081,172 peptides and 183,503 TCR pairs were split into training, validation, and test datasets according to the ratio of 0.8:0.1:0.1, respectively.

#### Peptide-MHC binding/presentation dataset

Binding affinity data between peptides and MHC was derived from the training data set of NetMHCpan-4.1^52^(https://services.healthtech.dtu.dk/suppl/immunology/NAR_NetMHCpan_NetMHCIIpan/), including 170,470 scaled and normalized half-maximal inhibitory concentration (IC_50_) values spanning 111 HLA class I alleles:

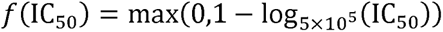

We randomly selected 10% of the affinity data for testing and trained the model on the remaining data since the original paper did not provide any independent test data. The training and validation data for predicting epitope presentation consisted of 288,032 eluted ligands (EL) and 16,739,285 negative pairs, respectively, across 149 alleles collected from BigMHC^53^ (https://data.mendeley.com/datasets/dvmz6pkzvb/4). The evaluation data comprising 45,409 ELs and 900,592 negative pairs spanning 36 alleles was the same EL dataset used in the NetMHCpan-4.1 study for benchmarking.

#### TCRαβ-pMHC recognition dataset

To construct a representative dataset for TCR-pMHC recognition, we combined the human TCR-pMHC binding pairs with confirmed CD8 expression from multiple sources, including IEDB^71^, VDJdb^56^, McPAS-TCR^80^, TBAdb^81^, 10X^82^, and Francis^66^. We associate the TCR sequences with the T-cell assays of specific epitope and MHC allele in IEDB (https://www.iedb.org/), which were downloaded on 31/08/2023 using the query parameters of Host: Homo sapiens, Reference type: journal article, linear epitope, MHC class I, and T cell assays only. We also downloaded the human TCR-pMHC-I datasets with paired TCRαβ sequences from VDJdb database (https://vdjdb.cdr3.net/), McPAS-TCR database (http://friedmanlab.weizmann.ac.il/McPAS-TCR/), and TBAdb from Pan immune repertoire database (PIRD, https://db.cngb.org/pird/), respectively. The 10X dataset was obtained from over 150,000 CD8^+^ T cells of four healthy donors staining with 44 distinct pMHC multimers. We integrated the binarized matrices and TCR clonotype annotations. We assigned the TCR binding specificities according to the criteria of UMI counts described in the application note “A new way of exploring immunity: linking highly multiplexed antigen recognition to immune repertoire and phenotype”. We also extracted the TCRαβ-pMHC binding pairs from CD8^+^ T cells of 28 SARS-CoV-2-infected patients and 23 unexposed individuals staining with SARS-CoV-2-derived DNA-barcoded-pMHC-multimers in the supplementary data file S3 of Francis et al., then removed the TCR clonotypes annotated with multiple alpha chains. We concatenated the TCRαβ-pMHC binding pairs containing CDR3αβ sequences, V-and J-gene annotations, peptide sequences, and MHC alleles from six original datasets into a combined dataset. The pre-processing of TCR-pMHC recognition data is presented in Supplementary Methods. Statistics of the datasets used for training and validation are shown in the **Supplementary Table 3**.

#### SARS-CoV-2 epitope-specific TCR clonotypes

The SARS-CoV-2-responsive TCR dataset was derived from a cohort of 55 individuals, including 16 SARS-CoV-2 negative participants, 30 participants recovered from mild disease, and 9 participants experienced symptomatic breakthrough infection that shaped spike-specific and non-spike-specific immune responses of memory CD8^+^ T cells upon infection and vaccination^45^. SARS-CoV-2 epitope-specific TCR clonotypes were identified and sequenced through DNA-barcoded MHC dextramers and single-cell TCR sequencing (scTCR-seq). This study assigned TCR recognition specificities for six spike protein epitopes and 12 non-spike epitopes presented on HLA alleles A*01:01, A*02:01, A*24:02, B*15:01, and B*44:02 according to the dextramer barcode UMI counts. We excluded two SARS-CoV-2 epitopes (A01_NTN and B44_VEN) from our analysis due to the minimal numbers of corresponding T cells and finally obtained 4471 TCR clonotypes. We removed the overlapped TCRαβ-pMHC pairs in our training dataset or the training data of MixTCRpred^30^. For external benchmarking, 3,540 TCR clonotypes with their experimentally assigned specificities were selected.

#### TCRαβ-pMHC complex structures

The crystal structures of the TCRαβ-pMHC complex were derived from the STCRDab^69^ (https://opig.stats.ox.ac.uk/webapps/stcrdab-stcrpred) database. After removing the noisy ones (PDB IDs: 6UZI, 7BYD) and duplicated TCRαβ-pMHC pairs, we constructed a structural dataset of 148 crystal structures. We extracted the coordinates of the heavy atoms of the amino acid residues and calculated the residue-level closest distances between CDR loops (CDR1α, CDR3α, and CDR3β) and the epitope. Contact residue pairs were defined as those whose spatial distances are within 5[, based on which the contact matrices were calculated and generated.

### Epitope-anchored contrastive transfer learning

#### Model backbone

Under the transfer learning paradigm, paired TCRαβ sequences of the binding or non-binding TCRs were sampled and input into the TCR language model to obtain the pre-trained TCR embeddings, respectively. At the same time, the representations for the pMHC complex were extracted from the peptide-MHC binding prediction model that took HLA molecules with their presented peptides as inputs. Model development of the pre-trained model can be found in Supplementary Methods. A multi-head self-attention layer and two 1x1 residual convolutional blocks were subsequently applied separately for further feature extraction from each sequence modality. Next, the fine-tuned embeddings of TCR and pMHC were fed into the contrastive co-embedding module or fused to provide model predictions for different downstream tasks.

#### Contrastive co-embedding module

The classification embeddings representing [CLS] tokens of TCR and pMHC were projected to a shared latent space by two MLP projectors. We designated one pMHC complex as an anchor in contrastive learning and then pulled the binding TCRs close to the anchor in the latent space while pushing the “non-binding” ones away. Given one pMHC complex *p*, a set of binding (positive) TCRs *T*_pos_, and a bunch of decoy (negative) TCRs *T*_neg_ with their projected representations in a training batch, cosine similarity between the pMHC anchor and sampled TCRs were calculated. The cosine similarities between TCR-pMHC binding pairs were expected to be larger than the similarities between the shuffled negative pairs. The epitope-anchored supervised contrastive loss^54^ was calculated as follows:

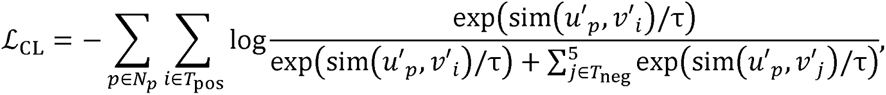

where sim(.) denotes cosine similarity, and *uʹ, vʹ*,’ represent the projected embeddings of pMHC and TCR, respectively. *τ* is the temperature factor of the loss function, *N_p_* is the collection of pMHC complexes in one batch, and five decoy TCRs are sampled each time.

#### Binding specificity prediction

We evaluated the model capacity to predict the binding specificities for unseen epitopes through five-fold cross-validation and assessed model generalizability on distinct TCR background populations from VDJdb. Epitopes in the training data were divided into groups by hierarchical clustering according to a minimum similarity score of 0.8 to achieve the zero-shot setting in cross-validation. The pairwise similarity score between epitope sequences *e_i_* and *e_j_* was defined as:

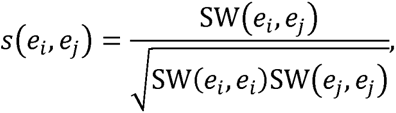

where SW(.) denotes the local alignment score between two protein sequences using the Smith-Waterman algorithm^83^ and BLOSUM62 substitution matrix. To predict TCR-pMHC binding specificities, classification [CLS] embeddings of TCR and pMHC were concatenated and input into an MLP classifier and sigmoid activation function. In addition to minimizing the contrastive loss, the binary cross entropy between predicted logits and labels was also included in the loss function to improve the adaptivity to unseen data.

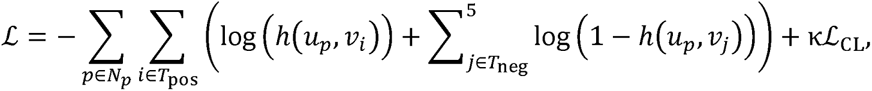

where *h*(*u,v*) denotes the predicted logits given the embeddings of pMHC and TCR, and *κ* is the weighting factor of the contrastive loss. Parameters of the pre-trained epitope language model, TCR language model, and MHC convolutional encoder were fixed. The cross-attention layer was fine-tuned to include TCR recognition information from MHC molecules. The AdamW optimizer with a learning rate of 2e-4 was used to train the binding specificity model for 50 epochs, and an early stopping strategy was employed to monitor the validation AUC.

#### Interaction conformation prediction

The residue-level interaction between CDR (CDR1α, CDR3α, CDR3β) sequences and epitope demonstrated an essential signature for the binding conformation of the TCR-pMHC complex. Thus, subsequently fed into a 2D convolutional layer with a kernel size of 3x3. The output of the the residue-level TCR and pMHC feature embeddings were integrated by outer product and convolutional layer consisted of two channels: the first channel was followed by a ReLU activation function to predict the pairwise distance matrices between CDRs and epitope; the second one used a sigmoid function to predict the contact probabilities between amino acid residue pairs. Five-fold cross-validation was performed in which highly similar epitopes were split into different folds (using the same strategy of epitope clustering in binding specificity prediction). A modified MSE loss divided by the distance between residues was utilized to reduce the influence on predictions from distant residue pairs, and binary cross-entropy loss was used for contact prediction. The weighting factors for interaction that involve CDR1α, CDR3α, and CDR3β were set to 0.3, 0.6, and 1.0, respectively, after taking into consideration of the sequence length and critical role of CDR3β. The two parts of loss were summed up and optimized using the AdamW optimizer with a learning rate of 2e-4 for 100 epochs. The pre-trained parameters were unfrozen in this stage, but the fine-tuning learning rate was ten times smaller.

All deep-learning models included in EPACT were implemented using PyTorch 2.0.1 and trained on one NVIDIA GeForce RTX 3090 GPU. Detailed model size and hyperparameters are provided in the **Supplementary Table 4**.

### Clustering analysis of epitope-specific TCR clones in co-embedding space

Representations of pMHC and TCR sequences were projected into the shared latent space, so we defined the embedding vector of a particular pMHC anchor as the centroid of the corresponding pMHC/epitope-specific TCR clusters. Therefore, candidate TCRs could be assigned to the closest pMHC anchor according to their cosine similarity. We also introduced a similarity threshold of 0.4 to maintain the high specificity of the epitope-specific TCR clusters. The pMHC anchors representing 16 SARS-CoV-2 epitopes and the epitope-specific TCR clones were visualized in two-dimensional space after Uniform Manifold Approximation and Projection^65^ (UMAP) with the parameter of n_neighbors=10, min_dist=0.1, and the metric is the cosine distance. We collected the CDR3αβ sequences in each SARS-CoV-2 epitope-specific TCR cluster, performed multiple sequence alignment (MSA) by MUSCLE^84^ and drew the CDR3 motifs, respectively. The positions in MSA where gaps occurred in over half of the aligned sequences were removed.

### Analysis of epitope-specific T-cell responses upon diverse SARS-CoV-2 infection and vaccination

As mentioned in the previous section, we predicted the SARS-CoV-2 epitope specificity of the TCR clonotypes according to the cosine distances to the pMHC anchors, thus constructing potential antigen-specific T cell clusters. After comparing the ratio of experimentally assigned epitope-specific TCRs in the one predicted cluster and others, we calculated the enrichment ratios in each cluster for each type of SARS-CoV-2 epitope-specific CD8+ T cells.

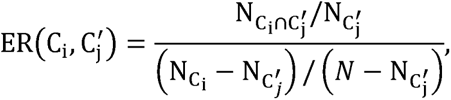

where C_i_, C_l_ represent the set of TCR clonotypes in experimental and predicted epitope-specific cluster *i*, and *j*, respectively, and *N* refers to the number of all clonotypes or in a particular cluster. We calculated the percentage prediction rank of TCRs to validate the relationship between T cell specificity and SARS-CoV-2 antigen exposure. Twenty thousand TCR sequences were sampled from the T cell repertoires of healthy human samples to generate the background distribution of binding scores, and we located the percentile for the candidate TCR. We also collected the expression profiles of various subsets of memory CD8^+^ T cells and metadata, including donors, vaccination category, and spike specificity from the original study^45^, to analyze the variation of binding specificity and clonal expansion upon diverse conditions.

### Calculation of contact scores and amino acid preference of peptides for cross-reactive TCRs

We predicted the residue-residue contact matrices between the cross-reactive AS-associated TCRs and their cognate peptides (viral peptides and self-peptides). The contact score of each amino acid residue along the peptide sequence was defined as the average of the top three contact probabilities with CDR1α or CDR3β residues. We also performed an in silico screening of cognate peptides for a particular TCR (MEL8/MEL5 TCR) by simulated annealing^85^ to investigate the consensus among binding peptides. Firstly, two thousand peptides were sampled from all HLA-A*02:01-presented epitopes deposited in the IEDB database as the initial peptide population. We predicted their binding scores with the target TCR and then randomly mutated a single amino acid of each peptide. After predicting the TCR binding specificity of the mutated sequences, the mutations with increased binding scores were accepted. In contrast, part of the other mutations was retained according to the acceptance probability.

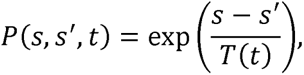

where *s* and *sʹ* denote the binding scores of the original and mutated peptide sequence, *T*(*t*) is the temperature of the *t*th iteration that declines proportionally. After 500 iterations, the top 2% of the final peptide population was extracted to render the sequence motif and heatmap representing the amino acid preferences of peptides for the cross-reactive TCRs.

#### Validation of interaction conformation between MEL8/MEL5 TCR and TAAs

We chose the TCR-pMHC complexes containing MEL8/MEL5 TCR and cognate tumor-associated antigens from the PDB database (PDB IDs: 7Q9A and 7Q9B) to validate the residue-level predictions of pairwise distances and contact probabilities. Contact residues from CDR loops and the epitope involved in van der Waals interactions (≤ 4 Å) and hydrogen bonds (≤ 3.4 Å) were selected for performance evaluation and visualized using PyMOL 2.4.0. We characterized the interaction conformations between MEL8/MEL5 TCR and all of the Melan A, BST2, and IMP2 peptides and compared them with the structural modeling results. The web server of TCRmodel2^86^ was employed to predict the 3D structures of TCR-pMHC complexes (modeling statistics in the **Supplementary Table 5**). We also computed the contact scores along CDR1α and CDR3β sequences with the HLA-A02-presented peptides that possibly bind to MEL8/MEL5 TCR (derived from binding specificity predictions by EPACT).

### Statistical analyses

All statistical tests in the study were two-sided. The error bars in the bar plots represent 95% confidence intervals unless otherwise stated. Performance benchmarking metrics, including AUC, AUPR, and RMSE, were calculated using the Python package scikit-learn 1.3.0. UMAP was performed using the Python package umap-learn 0.5.5. Local sequence alignment and hierarchical clustering of epitope sequences were performed using the Python package biopython 1.8.1 and scipy 1.11.1, respectively. Sequence motifs were visualized by the Python package logomaker 0.8 using the color scheme of “weblogo_protein”^87^. PyMOL 2.4.0 was used to visualize the 3D structure of TCR-pMHC complexes.

### Data availability

The curated datasets of TCR-pMHC recognition are shared and publicly accessible on GitHub https://github.com/zhangyumeng1sjtu/EPACT. Detailed information about the 10X Genomics Datasets is available in Table S2 and at https://www.10xgenomics.com/datasets. The crystal structures of TCR-pMHC complexes were obtained from the RCSB PDB database (https://www.rcsb.org/). Other structures listed in Table S5 were derived from TCRmodel2^86^

(https://tcrmodel.ibbr.umd.edu/) predictions. TCR sequences, experimental epitope specificity, gene expression, and other metadata of the SARS-CoV-2 responsive T cells were obtained from the original study^45^ (https://doi.org/10.1038/s41590-022-01184-4). Cross-reactive TCRs and activated peptides in the context of HLA-B*27:05 were obtained from the original study^46^ (https://doi.org/10.1038/s41586-022-05501-7). Binding hotspots between MEL8 or MEL5 TCR and corresponding pMHC complexes were derived from the original study^47^ (https://doi.org/10.1016/j.cell.2023.06.020).

### Code availability

The source code and model weights of EPACT are available on GitHub https://github.com/zhangyumeng1sjtu/EPACT.

---

**Extended Data Fig. 1.**
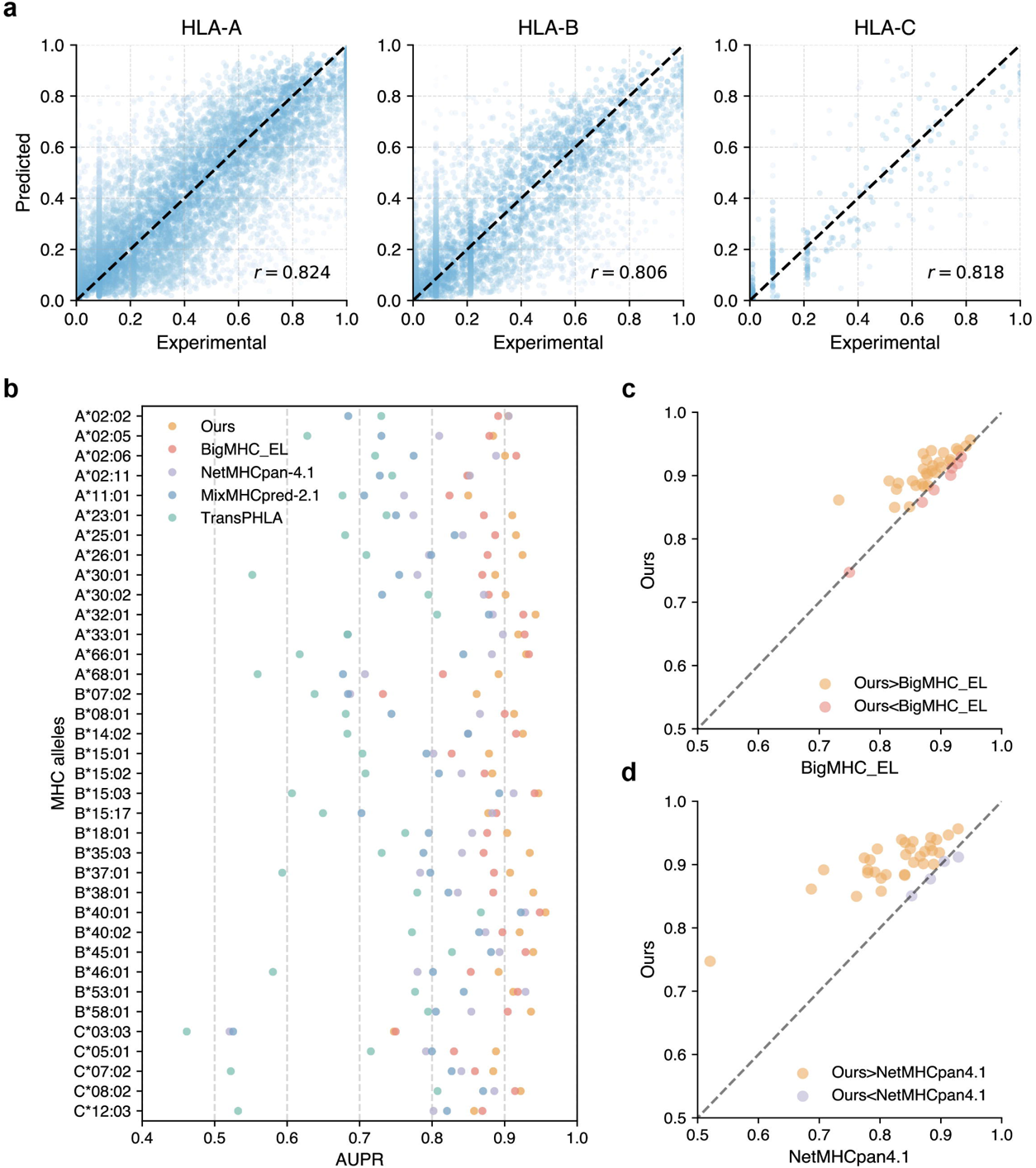
Performance of pMHC binding affinity and eluted ligand model. **a,** The experimental and predicted binding affinity (normalized IC_50_ values) of tested peptide-MHC pairs by stratifying HLA subtypes. **b,** Performance of our epitope presentation model and existing methods (BigMHC, NetMHCpan-4.1, MixMHCpred-2.1, and TransPHLA) on each test HLA molecule. Pairwise comparisons between the predicted AUPR of our epitope presentation model and **c,** BigMHC and **d,** NetMHCpan-4. Each point represents one HLA molecule, and the orange point indicates the better performance of our model.

**Extended Data Fig. 2.**
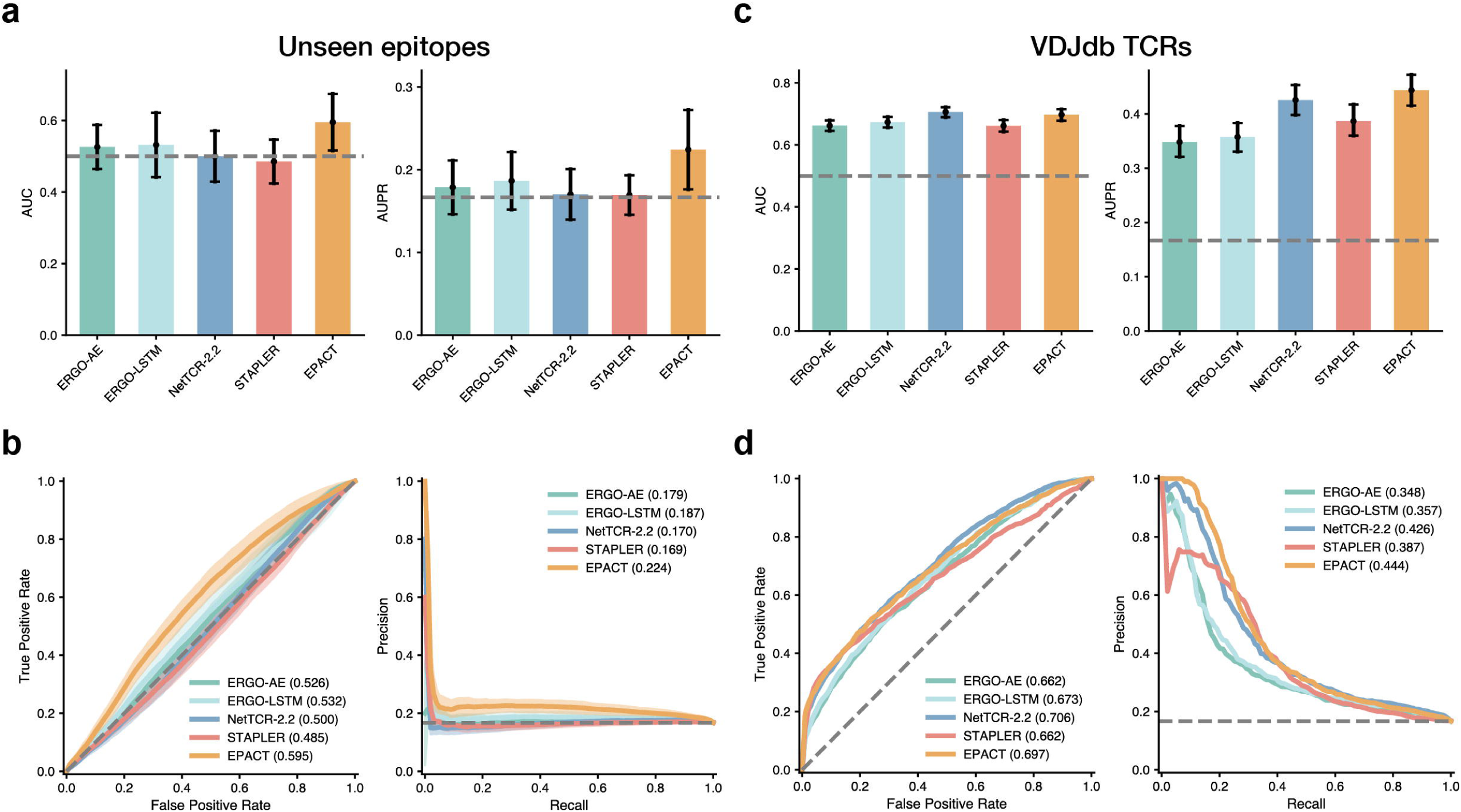
Benchmarking results on paired TCRαβ binding specificity data. **a,** Bar plots of AUCs and AUPRs, and **b,** ROC curves and precision-recall curves of the candidate TCRαβ models in cross-validation (i.e., prediction of unseen epitopes). The error bars indicate the standard deviation across five-folds, and the shade around the curves represents the standard error of TPRs and precisions. **c,** Bar plots of AUCs and AUPRs, and **d,** ROC curves and precision-recall curves of the candidate TCRαβ models in testing (predicting for VDJdb TCRs). The error bars indicate the 95% confidence intervals by 1000 bootstrap iterations. The gray dashed lines denote the results of random predictions.

**Extended Data Fig. 3.**
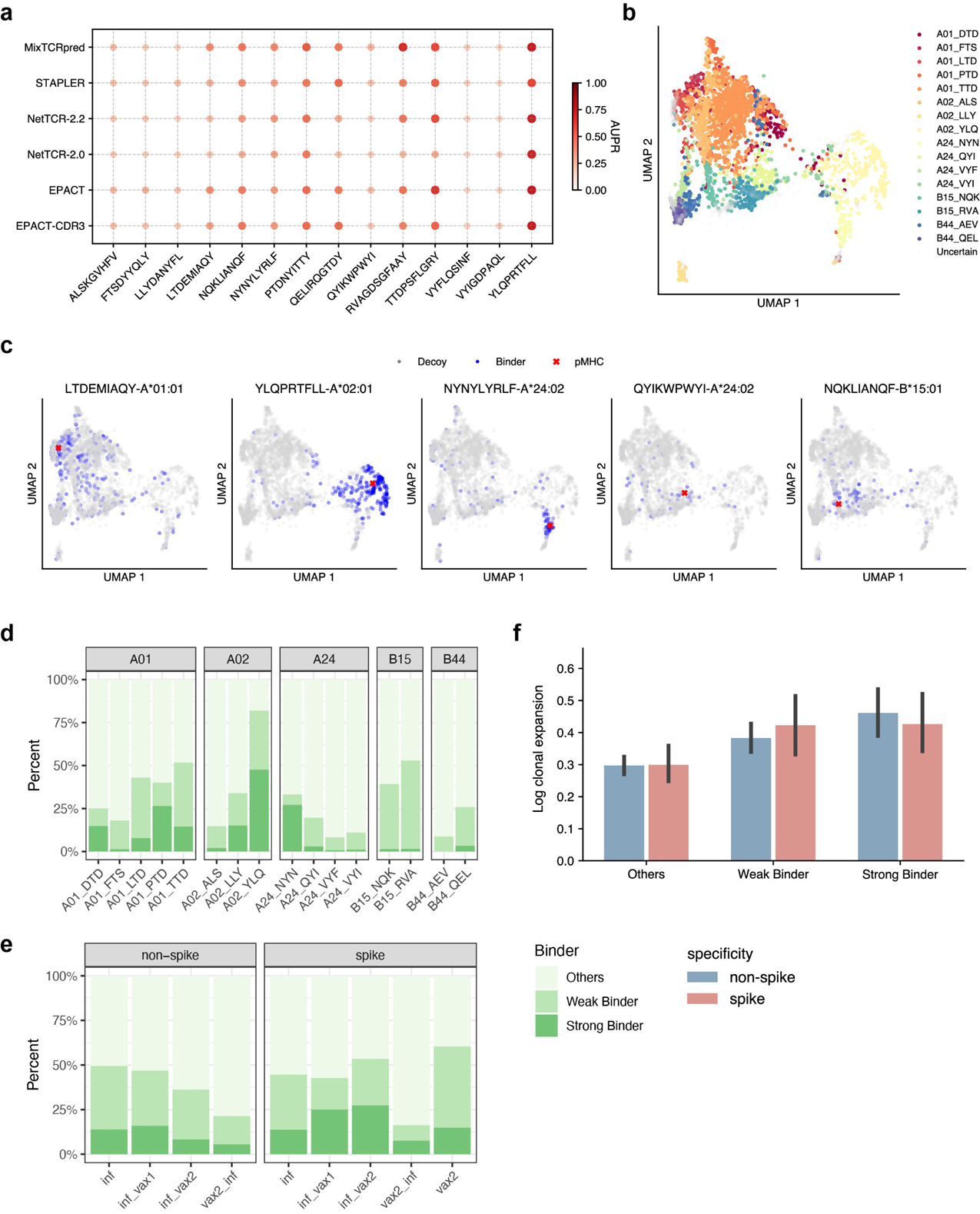
Interpretable prediction and analysis of SARS-CoV-2-responsive TCR clonotypes. **a,** Performance comparison in terms of AUPRs derived from MixTCRpred, STAPLER, NetTCR-2.0, NetTCR-2.2, and EPACT for 14 SARS-CoV-2 epitopes. The darker color and larger size of the point indicate a higher AUPR. **b,** UMAP projection of the predicted SARS-CoV-2 epitope-specific TCR clusters in the unseen SARS-CoV-2-responsive TCR dataset. **c,** UMAP projections of five spike epitope targets and experimental binding TCRs. The red cross in each subplot denotes the pMHC anchor, the blue points represent the binding TCRs, and the gray points indicate the decoys TCRs with no experimental evidence responding to the target. Proportions of predicted strong binders (rank≥99.5%) and weak binders (rank≥95%) **d,** targeting each SARS-CoV-2 epitope and **e.** in spike-specific or non-spike-specific TCRs across different categories of SARS-CoV-2 infection and vaccination. **f.** Bar plots showing the log clonal expansion of the strong and weak TCR binders.

**Extended Data Fig. 4.**
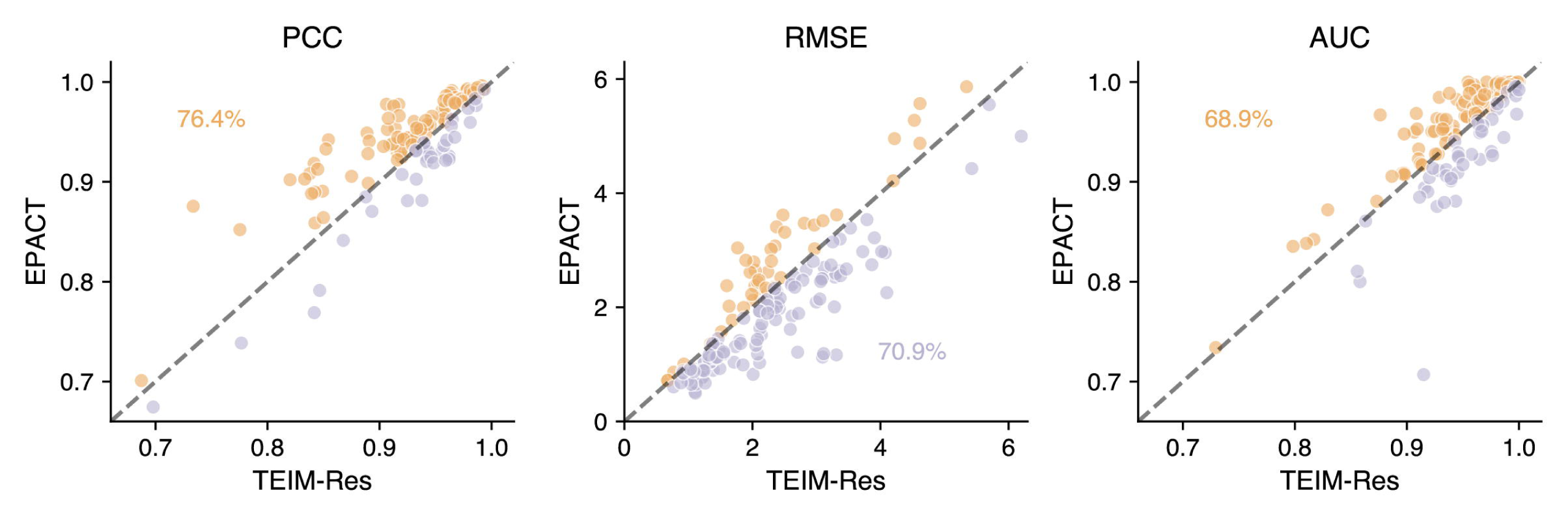
Pairwise comparison of predicted CDR3β-epitope interactions by EPACT and TEIM-Res. Scatter plots displaying the validation PCC (left), RMSE (middle), and AUC (right) predicted by EPACT and TEIM-Res for CDR3β-epitope interactions in each TCR-pMHC crystal structure. The points colored in orange and purple indicate the values of EPACT’s metrics are higher or lower than those predicted by TEIM-Res, respectively.

**Extended Data Fig. 5.**
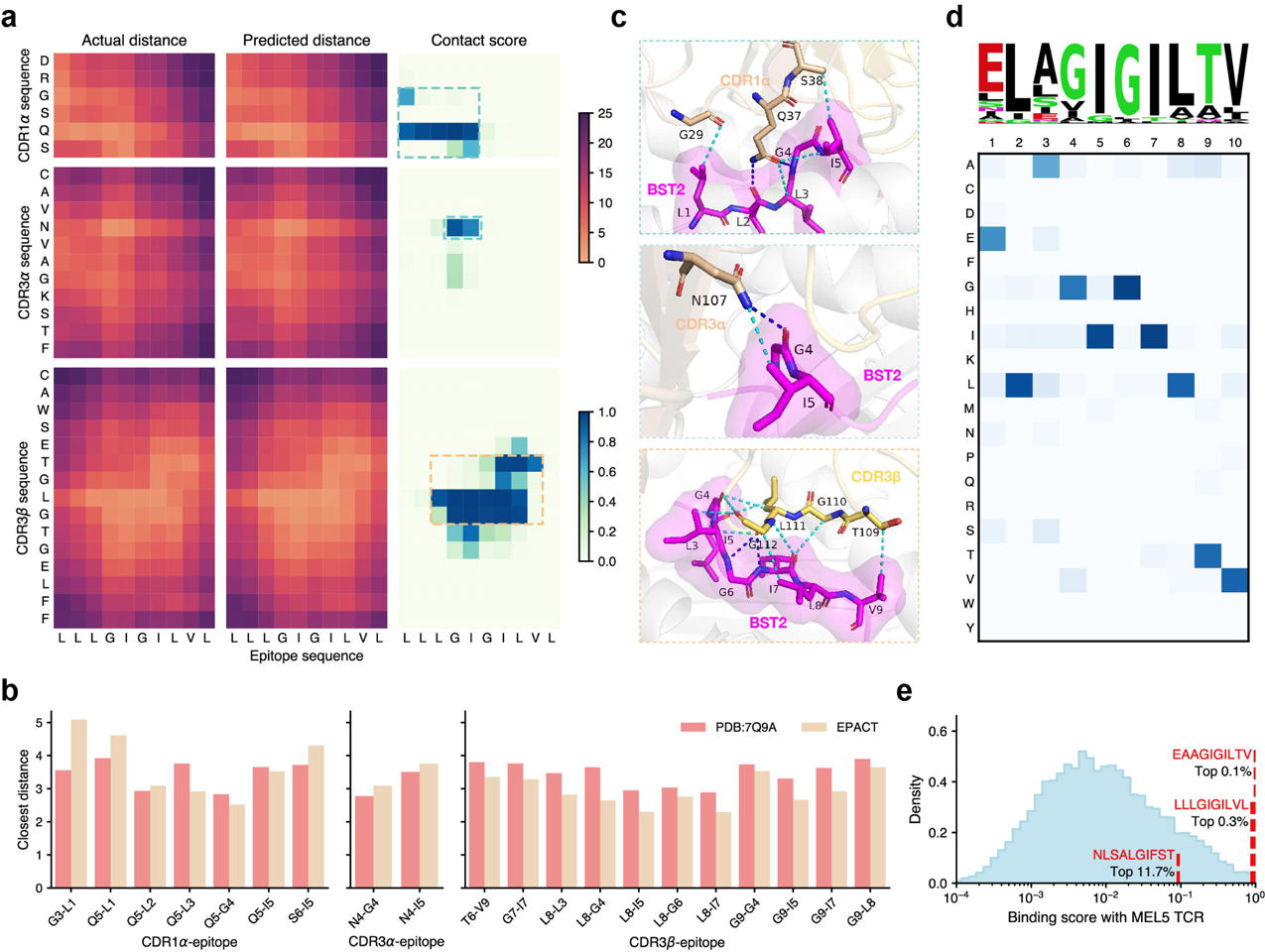
EPACT identifies the recognition between three tumor-associated epitopes and MEL5 TCR. **a,** Residue-residue experimental (left) and predicted (middle) distance matrices and predicted contact scores (right) characterizing CDR1α-epitope (top), CDR3α-epitope (middle), and CDR3β-epitope (bottom) interactions in the MEL5 TCR-Melan A peptide-HLA-A*02:01 complex. The predicted interactions were derived from validation test. The color scales in the heatmap represent amino acid pairs from close to distant and contact scores from low to high. The core interaction regions are surrounded by the dashed lines. **b,** Bar plots to compare the experimental distances in PDB structures (PDB: 7Q9A) and predicted distances by EPACT of the inter-chain contact residue pairs (≤4Å) from CDR1α/CDR3α/CDR3β and Melan A peptide. **c,** Visualization of the core interaction regions, including CDR1α (left), CDR3α (middle), and CDR3β (right) loops of MEL5 TCR and Melan A peptide. The cyan and dark blue line between residues denote van der Waals interaction and hydrogen bond, respectively. **d,** Sequence motif (top) and heatmap (bottom) to display the positional amino acid preferences of peptides recognized by MEL5 TCR. **e,** Density plots showing the distribution of predicted binding scores to MEL5 TCR among the IEDB HLA-A*02:01-presented peptides. The x-axis is transformed into a log scale.

**Supplementary Fig. 1.** Detailed model architecture of EPACT.

**Supplementary Fig. 2.** Epitope-level performance in predicting TCR binding to unseen peptides.

**Supplementary Fig. 3.** Evaluation of model robustness under various peptide lengths and similarities.

**Supplementary Fig. 4.** CDR3 motifs of spike-epitope-specific TCRαβ sequences.

**Supplementary Fig. 5.** Co-embedding visualization of SARS-CoV-2 antigen(non-spike)-specific TCRs.

**Supplementary Fig. 6.** EPACT recognizes the interacting residues between the MEL8 CDR1α sequence and cognate epitopes.

**Supplementary Fig. 7.** Mutual corroboration of EPACT-predicted interactions and structure modeling of MEL8 TCR-pMHC complex.

**Supplementary Table 1.** Benchmarking results and ablation study of EPACT.

**Supplementary Table 2.** 10X Genomics datasets used for pre-training human TCRs.

**Supplementary Table 3.** Statistics of datasets used for model training.

**Supplementary Table 4.** Statistics of model size and hyperparameters.

**Supplementary Table 5.** Statistics of TCRmodel2-predicted TCR-pMHC complex structures.

## Notes

### Competing Interest Statement

The authors have declared no competing interest.

## References

1. Zhang, N. & Bevan, Michael J. CD8+ T Cells: Foot Soldiers of the Immune System. Immunity 35, 161–168 (2011).

2. Petrelli, A. & van Wijk, F. CD8+ T cells in human autoimmune arthritis: the unusual suspects. Nature Reviews Rheumatology 12, 421–428 (2016).

3. Reina-Campos, M., Scharping, N.E. & Goldrath, A.W. CD8+ T cell metabolism in infection and cancer. Nature Reviews Immunology 21, 718–738 (2021).

4. Raskov, H., Orhan, A., Christensen, J.P. & Gögenur, I. Cytotoxic CD8+ T cells in cancer and cancer immunotherapy. British Journal of Cancer 124, 359–367 (2021).

5. Philip, M. & Schietinger, A. CD8+ T cell differentiation and dysfunction in cancer. Nat Rev Immunol 22, 209–223 (2022).

6. Tian, S., Maile, R., Collins, E.J. & Frelinger, J.A. CD8+ T Cell Activation Is Governed by TCR-Peptide/MHC Affinity, Not Dissociation Rate. J Immun 179, 2952–2960 (2007).

7. Rossjohn, J. et al. T Cell Antigen Receptor Recognition of Antigen-Presenting Molecules. Annu Rev Immunol 33, 169–200 (2015).

8. Chandran, S.S. & Klebanoff, C.A. T cell receptor-based cancer immunotherapy: Emerging efficacy and pathways of resistance. Immunological Reviews 290, 127–147 (2019).

9. Cowell, L.G. The Diagnostic, Prognostic, and Therapeutic Potential of Adaptive Immune Receptor Repertoire Profiling in Cancer. Cancer Res 80, 643–654 (2020).

10. Kidman, J. et al. Characteristics of TCR repertoire associated with successful immune checkpoint therapy responses. Front Immunol 11, 587014 (2020).

11. Valpione, S. et al. The T cell receptor repertoire of tumor infiltrating T cells is predictive and prognostic for cancer survival. Nat Commun 12, 4098 (2021).

12. Pai, J.A. & Satpathy, A.T. High-throughput and single-cell T cell receptor sequencing technologies. Nature Methods 18, 881–892 (2021).

13. Zhang, S.-Q. et al. High-throughput determination of the antigen specificities of T cell receptors in single cells. Nature Biotechnology 36, 1156–1159 (2018).

14. Ng, A.H.C. et al. MATE-Seq: microfluidic antigen-TCR engagement sequencing. Lab Chip 19, 3011–3021 (2019).

15. Sewell, A.K. Why must T cells be cross-reactive? Nat Rev Immunol 12, 669–677 (2012).

16. Spear, T.T., Evavold, B.D., Baker, B.M. & Nishimura, M.I. Understanding TCR affinity, antigen specificity, and cross-reactivity to improve TCR gene-modified T cells for cancer immunotherapy. Cancer Immunol Immunother 68, 1881–1889 (2019).

17. Cole, D.K. et al. Hotspot autoimmune T cell receptor binding underlies pathogen and insulin peptide cross-reactivity. J Clin Invest 126, 2191–2204 (2016).

18. Cusick, M.F., Libbey, J.E. & Fujinami, R.S. Molecular Mimicry as a Mechanism of Autoimmune Disease. Clin Rev Allergy Immunol 42, 102–111 (2012).

19. Linette, G.P. et al. Cardiovascular toxicity and titin cross-reactivity of affinity-enhanced T cells in myeloma and melanoma. Blood 122, 863–871 (2013).

20. Szeto, C., Lobos, C.A., Nguyen, A.T. & Gras, S. TCR Recognition of Peptide–MHC-I: Rule Makers and Breakers. Int J Mol Sci 22, 68 (2021).

21. Hudson, D., Fernandes, R.A., Basham, M., Ogg, G. & Koohy, H. Can we predict T cell specificity with digital biology and machine learning? Nature Reviews Immunology 23, 511–521 (2023).

22. Huang, H., Wang, C., Rubelt, F., Scriba, T.J. & Davis, M.M. Analyzing the Mycobacterium tuberculosis immune response by T-cell receptor clustering with GLIPH2 and genome-wide antigen screening. Nat Biotechnol 38, 1194–1202 (2020).

23. Sidhom, J.-W., Larman, H.B., Pardoll, D.M. & Baras, A.S. DeepTCR is a deep learning framework for revealing sequence concepts within T-cell repertoires. Nat Commun 12, 1605 (2021).

24. Mayer-Blackwell, K. et al. TCR meta-clonotypes for biomarker discovery with tcrdist3 enabled identification of public, HLA-restricted clusters of SARS-CoV-2 TCRs. eLife 10, e68605 (2021).

25. Kevin, W., et al. TCR-BERT: learning the grammar of T-cell receptors for flexible antigen-xbinding analyses. bioRxiv, 2021.2011.2018.469186 (2021).

26. Gielis, S. et al. Detection of enriched T cell epitope specificity in full T cell receptor sequence repertoires. Front Immunol 10, 2820 (2019).

27. Jokinen, E., Huuhtanen, J., Mustjoki, S., Heinonen, M. & Lähdesmäki, H. Predicting recognition between T cell receptors and epitopes with TCRGP. PLoS Comput Biol 17, e1008814 (2021).

28. Montemurro, A. et al. NetTCR-2.0 enables accurate prediction of TCR-peptide binding by using paired TCRα and β sequence data. Commun Biol 4, 1060 (2021).

29. Zhang, W. et al. A framework for highly multiplexed dextramer mapping and prediction of T cell receptor sequences to antigen specificity. Sci Adv 7, eabf5835 (2021).

30. Giancarlo, C. et al. Deep learning predictions of TCR-epitope interactions reveal epitope-specific chains in dual alpha T cells. bioRxiv, 2023.2009.2013.557561 (2023).

31. Springer, I., Tickotsky, N. & Louzoun, Y. Contribution of T cell receptor alpha and beta CDR3, MHC typing, V and J genes to peptide binding prediction. Front Immunol 12, 664514 (2021).

32. Weber, A., Born, J. & Rodriguez Martínez, M. TITAN: T-cell receptor specificity prediction with bimodal attention networks. Bioinformatics 37, i237–i244 (2021).

33. Lu, T. et al. Deep learning-based prediction of the T cell receptor–antigen binding specificity. Nat Mach Intell 3, 864–875 (2021).

34. Peng, X. et al. Characterizing the interaction conformation between T-cell receptors and epitopes with deep learning. Nat Mach Intell 5, 395–407 (2023).

35. Gao, Y. et al. Pan-Peptide Meta Learning for T-cell receptor–antigen binding recognition. Nat Mach Intell 5, 236–249 (2023).

36. Bjørn, P.Y.K., et al. STAPLER: Efficient learning of TCR-peptide specificity prediction from full-length TCR-peptide data. bioRxiv, 2023.2004.2025.538237 (2023).

37. Ethan, F., Manjima, D. & Binbin, C. TAPIR: a T-cell receptor language model for predicting rare and novel targets. bioRxiv, 2023.2009.2012.557285 (2023).

38. Barthelemy, M.-P., Christoph, F., Martin, W., Aleksandra, M.W. & Thierry, M. TULIP — a Transformer based Unsupervised Language model for Interacting Peptides and T-cell receptors that generalizes to unseen epitopes. bioRxiv, 2023.2007.2019.549669 (2024).

39. Jensen, M.F. & Nielsen, M. NetTCR 2.2 - Improved TCR specificity predictions by combining pan- and peptide-specific training strategies, loss-scaling and integration of sequence similarity. eLife 12, RP93934 (2023).

40. Yi, H. et al. pan-MHC and cross-Species Prediction of T Cell Receptor-Antigen Binding. bioRxiv, 2023.2012.2001.569599 (2023).

41. Spindler, M.J. et al. Massively parallel interrogation and mining of natively paired human TCRαβ repertoires. Nature Biotechnology 38, 609–619 (2020).

42. Dens, C., Laukens, K., Bittremieux, W. & Meysman, P. The pitfalls of negative data bias for the T-cell epitope specificity challenge. Nature Machine Intelligence 5, 1060–1062 (2023).

43. Song, I. et al. Broad TCR repertoire and diverse structural solutions for recognition of an immunodominant CD8+ T cell epitope. Nat Struct Mol Biol 24, 395–406 (2017).

44. Singh, R., Sledzieski, S., Bryson, B., Cowen, L. & Berger, B. Contrastive learning in protein language space predicts interactions between drugs and protein targets. Proc Natl Acad Sci U S A 120, e2220778120 (2023).

45. Minervina, A.A. et al. SARS-CoV-2 antigen exposure history shapes phenotypes and specificity of memory CD8+ T cells. Nat Immunol 23, 781–790 (2022).

46. Yang, X. et al. Autoimmunity-associated T cell receptors recognize HLA-B*27-bound peptides. Nature 612, 771–777 (2022).

47. Dolton, G. et al. Targeting of multiple tumor-associated antigens by individual T cell receptors during successful cancer immunotherapy. Cell 186, 3333–3349.e3327 (2023).

48. Bepler, T. & Berger, B. Learning the protein language: Evolution, structure, and function. Cell Syst 12, 654–669.e653 (2021).

49. Atchley, W.R., Zhao, J., Fernandes, A.D. & Drüke, T. Solving the protein sequence metric problem. Proc Natl Acad Sci U S A 102, 6395–6400 (2005).

50. He, K., Zhang, X., Ren, S. & Sun, J. in Proceedings of the IEEE conference on computer vision and pattern recognition 770–778 (2016).

51. Neefjes, J., Jongsma, M.L.M., Paul, P. & Bakke, O. Towards a systems understanding of MHC class I and MHC class II antigen presentation. Nat Rev Immunol 11, 823–836 (2011).

52. Reynisson, B., Alvarez, B., Paul, S., Peters, B. & Nielsen, M. NetMHCpan-4.1 and NetMHCIIpan-4.0: improved predictions of MHC antigen presentation by concurrent motif deconvolution and integration of MS MHC eluted ligand data. Nucleic Acids Res 48, W449–W454 (2020).

53. Albert, B.A. et al. Deep neural networks predict class I major histocompatibility complex epitope presentation and transfer learn neoepitope immunogenicity. Nat Mach Intell 5, 861–872 (2023).

54. Khosla, P. et al. in Advances in neural information processing systems, Vol. 33 18661–18673 (2020).

55. Sethna, Z. et al. Population variability in the generation and selection of T-cell repertoires. PLoS Comput Biol 16, e1008394 (2020).

56. Goncharov, M. et al. VDJdb in the pandemic era: a compendium of T cell receptors specific for SARS-CoV-2. Nat Methods 19, 1017–1019 (2022).

57. Feng, D., Bond, C.J., Ely, L.K., Maynard, J. & Garcia, K.C. Structural evidence for a germline-encoded T cell receptor–major histocompatibility complex interaction ’codon’. Nat Immunol 8, 975–983 (2007).

58. Meysman, P. et al. Benchmarking solutions to the T-cell receptor epitope prediction problem: IMMREP22 workshop report. ImmunoInformatics 9, 100024 (2023).

59. San, D. et al. Structural basis of the TCR-pHLA complex provides insights into the unconventional recognition of CDR3β in TCR cross-reactivity and alloreactivity. Cell Insight 2, 100076 (2023).

60. Lefranc, M.-P. et al. IMGT unique numbering for immunoglobulin and T cell receptor variable domains and Ig superfamily V-like domains. Dev Comp Immunol 27, 55–77 (2003).

61. Borràs, D.M. et al. Single cell dynamics of tumor specificity vs bystander activity in CD8+ T cells define the diverse immune landscapes in colorectal cancer. Cell Discov 9, 114 (2023).

62. Zhang, B. et al. Multimodal single-cell datasets characterize antigen-specific CD8+ T cells across SARS-CoV-2 vaccination and infection. Nat Immunol 24, 1725–1734 (2023).

63. Huuhtanen, J. et al. Evolution and modulation of antigen-specific T cell responses in melanoma patients. Nat Commun 13, 5988 (2022).

64. Reiser, J.-B. et al. CDR3 loop flexibility contributes to the degeneracy of TCR recognition. Nat Immunol 4, 241–247 (2003).

65. McInnes, L., Healy, J. & Melville, J. Umap: Uniform manifold approximation and projection for dimension reduction. Preprint at 10.48550/arXiv.41802.03426 (2018).

66. Francis, J.M., et al. Allelic variation in class I HLA determines CD8+ T cell repertoire shape and cross-reactive memory responses to SARS-CoV-2. Sci Immunol 7, eabk3070 (2022).

67. Shomuradova, A.S. et al. SARS-CoV-2 Epitopes Are Recognized by a Public and Diverse Repertoire of Human T Cell Receptors. Immunity 53, 1245–1257.e1245 (2020).

68. van der Leun, A.M., Thommen, D.S. & Schumacher, T.N. CD8+ T cell states in human cancer: insights from single-cell analysis. Nat Rev Cancer 20, 218–232 (2020).

69. Leem, J., de Oliveira, S.H P., Krawczyk, K. & Deane, C.M. STCRDab: the structural T-cell receptor database. Nucleic Acids Res 46, D406–D412 (2017).

70. Andersen, R. et al. Long-Lasting Complete Responses in Patients with Metastatic Melanoma after Adoptive Cell Therapy with Tumor-Infiltrating Lymphocytes and an Attenuated IL2 Regimen. Clin Cancer Res 22, 3734–3745 (2016).

71. Vita, R. et al. The Immune Epitope Database (IEDB): 2018 update. Nucleic Acids Res 47, D339–D343 (2018).

72. Wooldridge, L. et al. A Single Autoimmune T Cell Receptor Recognizes More Than a Million Different Peptides. J Biol Chem 287, 1168–1177 (2012).

73. Birnbaum, Michael E. et al. Deconstructing the Peptide-MHC Specificity of T Cell Recognition. Cell 157, 1073–1087 (2014).

74. Bradley, P. Structure-based prediction of T cell receptor:peptide-MHC interactions. eLife 12, e82813 (2023).

75. Yost, K.E. et al. Clonal replacement of tumor-specific T cells following PD-1 blockade. Nat Med 25, 1251–1259 (2019).

76. Kourtis, N. et al. A single-cell map of dynamic chromatin landscapes of immune cells in renal cell carcinoma. Nat Cancer 3, 885–898 (2022).

77. Liu, T. et al. Single cell profiling of primary and paired metastatic lymph node tumors in breast cancer patients. Nat Commun 13, 6823 (2022).

78. Zheng, L. et al. Pan-cancer single-cell landscape of tumor-infiltrating T cells. Science 374, abe6474 (2021).

79. Wu, T.D. et al. Peripheral T cell expansion predicts tumour infiltration and clinical response. Nature 579, 274–278 (2020).

80. Tickotsky, N., Sagiv, T., Prilusky, J., Shifrut, E. & Friedman, N. McPAS-TCR: a manually curated catalogue of pathology-associated T cell receptor sequences. Bioinformatics 33, 2924–2929 (2017).

81. Zhang, W. et al. PIRD: Pan Immune Repertoire Database. Bioinformatics 36, 897–903 (2019).

82. Genomics, x. A new way of exploring immunity–linking highly multiplexed antigen recognition to immune repertoire and phenotype. 10x Genomics. (2019).

83. Smith, T.F. & Waterman, M.S. Identification of common molecular subsequences. J Mol Biol 147, 195–197 (1981).

84. Edgar, R.C. MUSCLE: multiple sequence alignment with high accuracy and high throughput. Nucleic Acids Res 32, 1792–1797 (2004).

85. Kirkpatrick, S., Gelatt, C.D. & Vecchi, M.P. Optimization by Simulated Annealing. Science 220, 671–680 (1983).

86. Yin, R. et al. TCRmodel2: high-resolution modeling of T cell receptor recognition using deep learning. Nucleic Acids Res 51, W569–W576 (2023).

87. Tareen, A. & Kinney, J.B. Logomaker: beautiful sequence logos in Python. Bioinformatics 36, 2272–2274 (2019).

