## Supplementary material for "Epitope-anchored contrastive transfer learning for paired CD8^+^ cell receptor-antigen recognition": Supplenmetary figures and methods

Supplementary figures

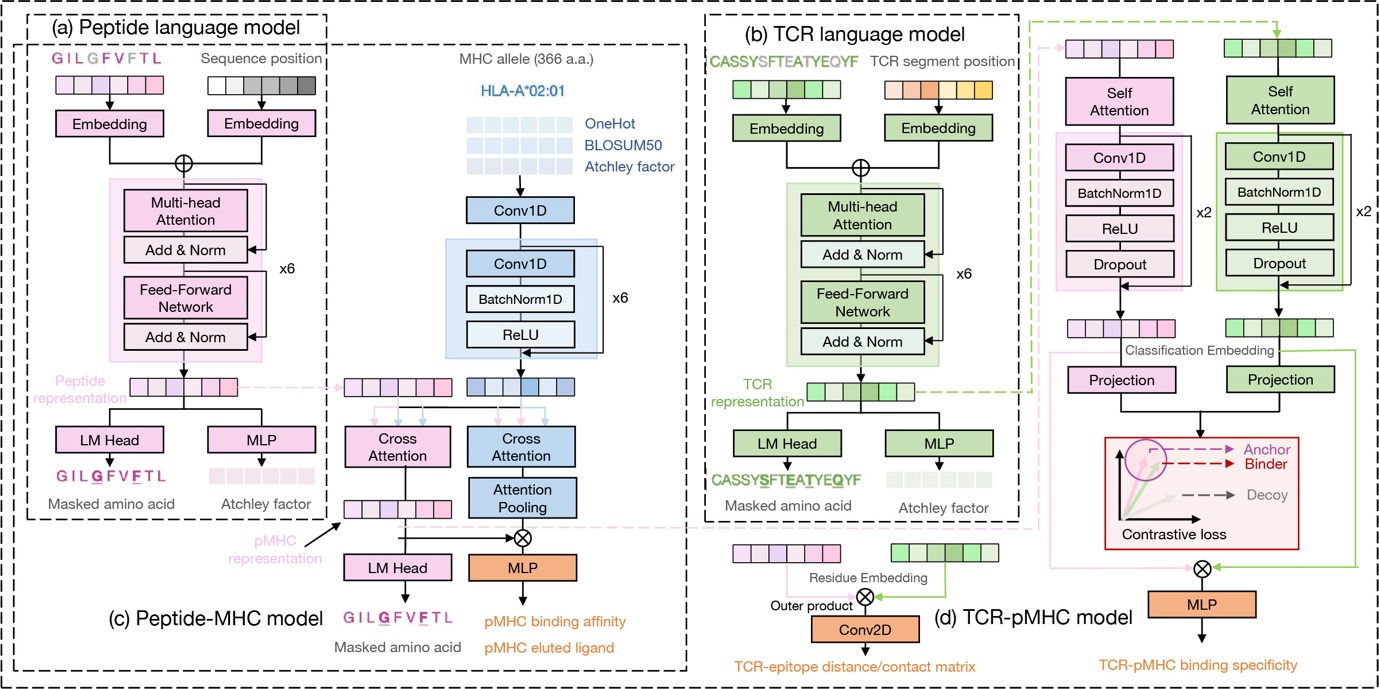

**Supplementary Fig. 1** Detailed model architecture of EPACT.

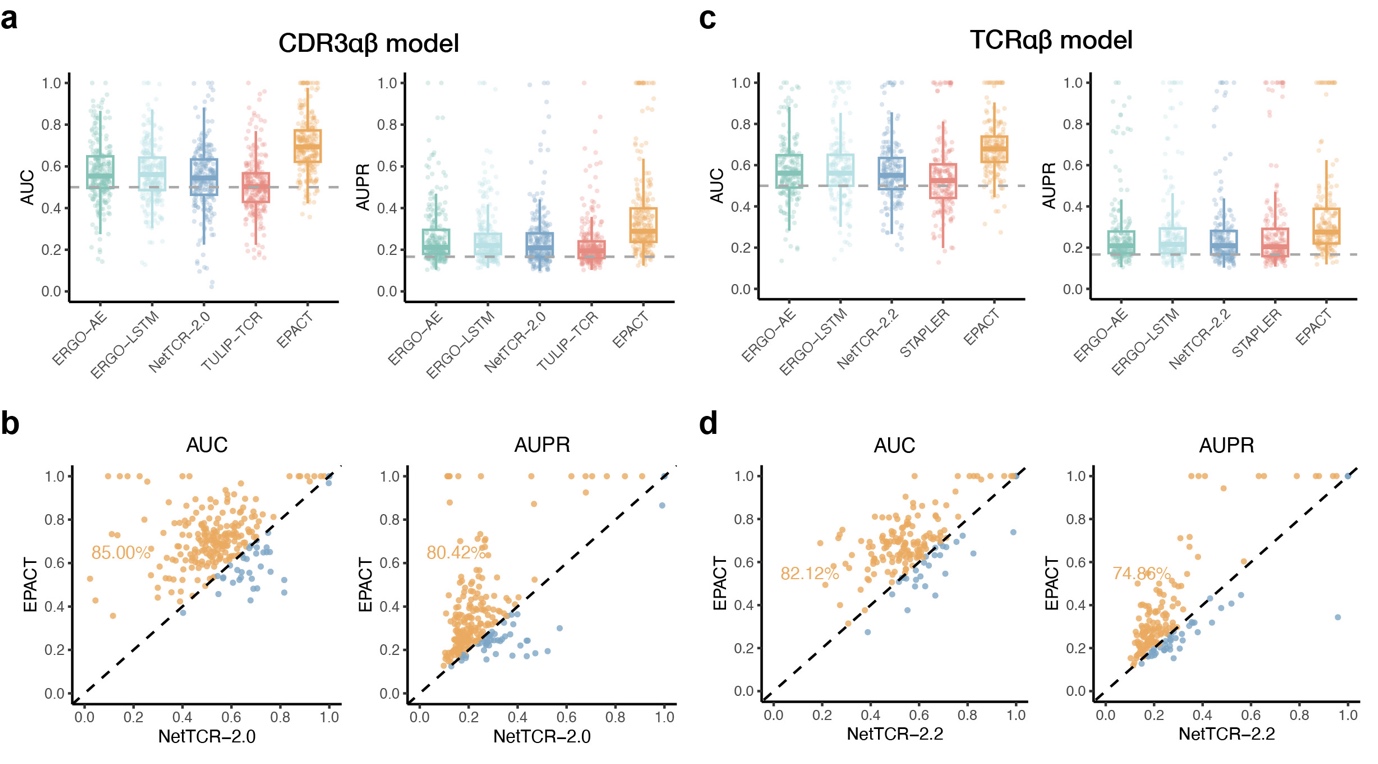

**Supplementary Fig. 2** Epitope-level performance in predicting TCR binding to unseen peptides.

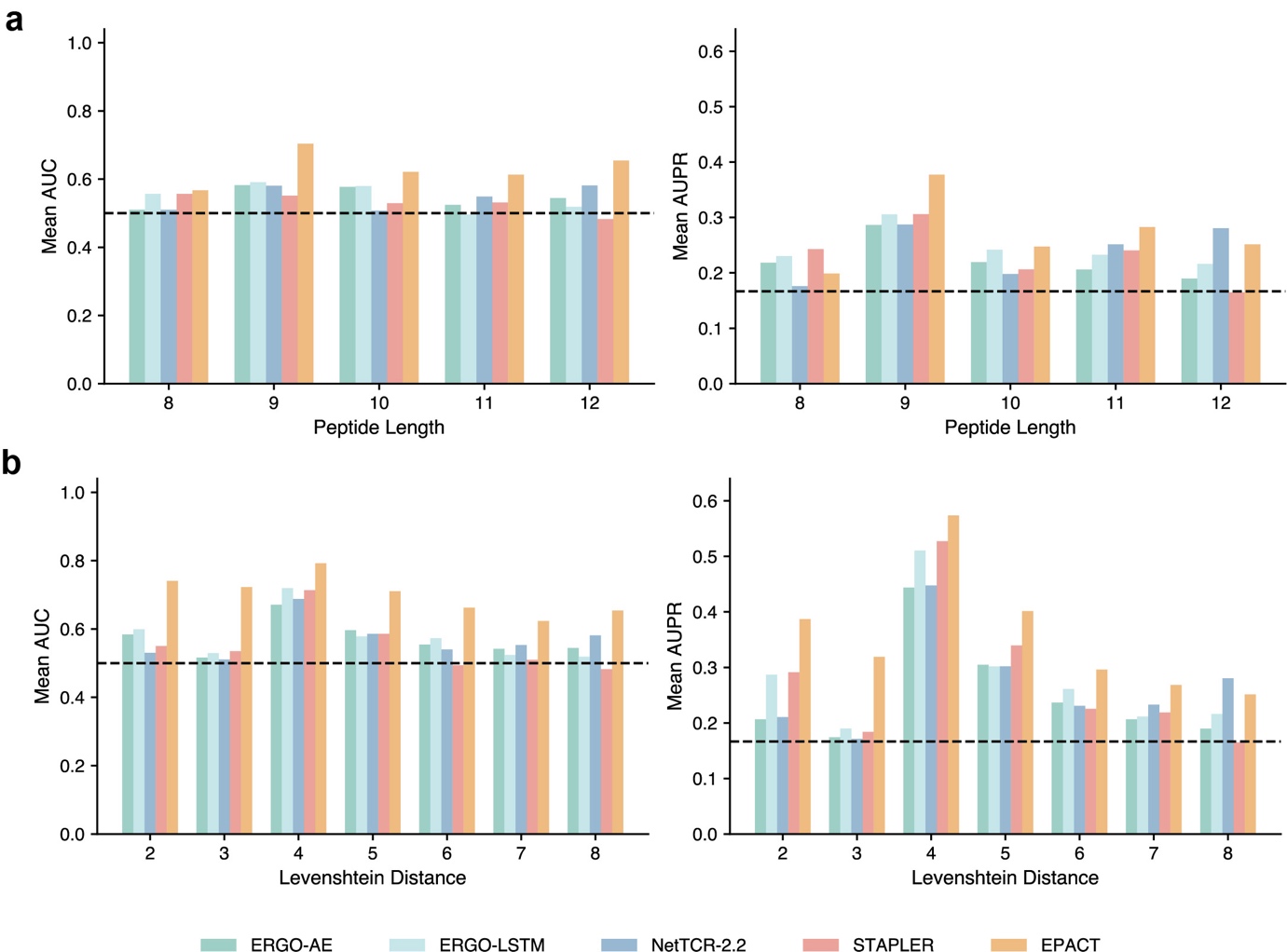

**Supplementary Fig. 3** Evaluation of model robustness under various peptide lengths and similarities.

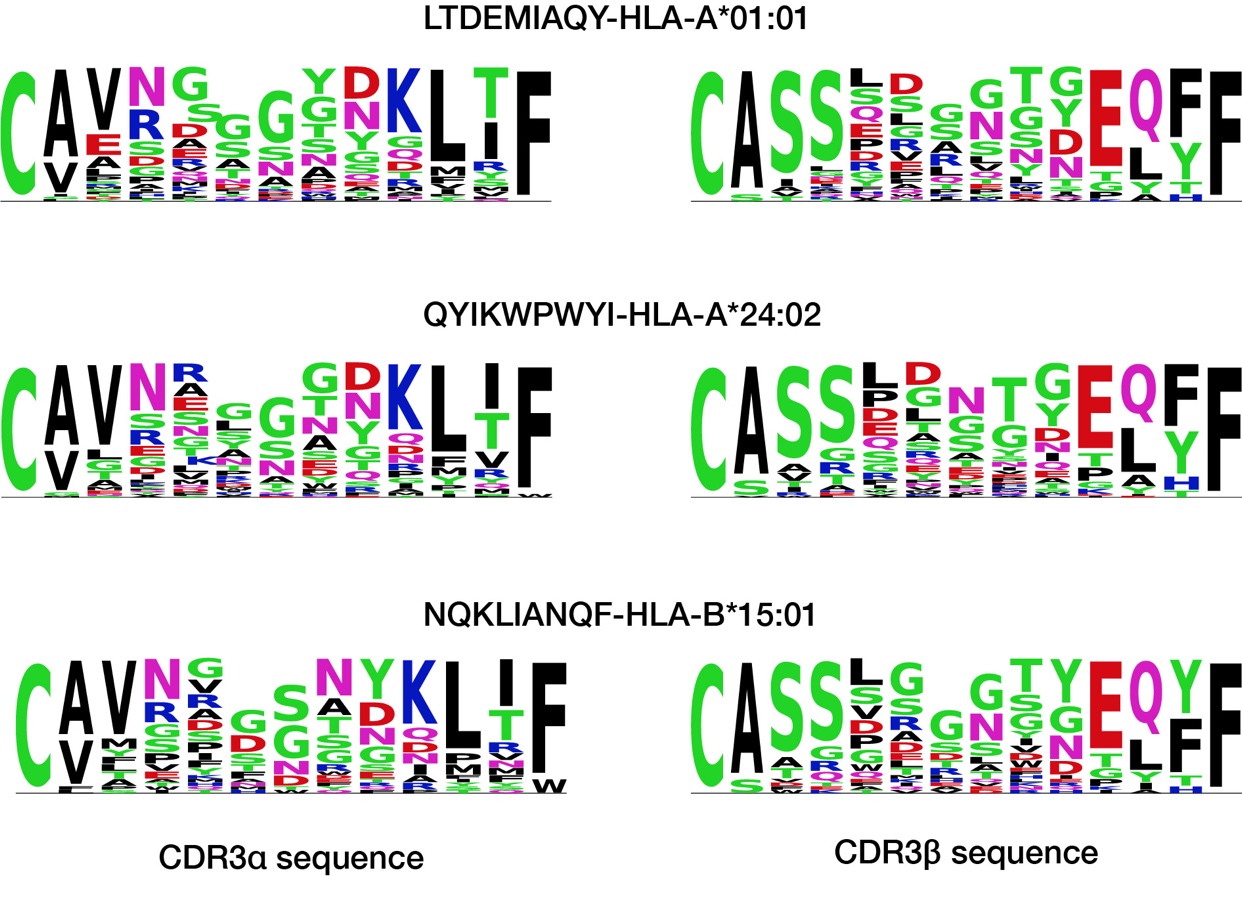

**Supplementary Fig. 4** CDR3 motifs of spike-epitope-specific TCRαβ sequences.

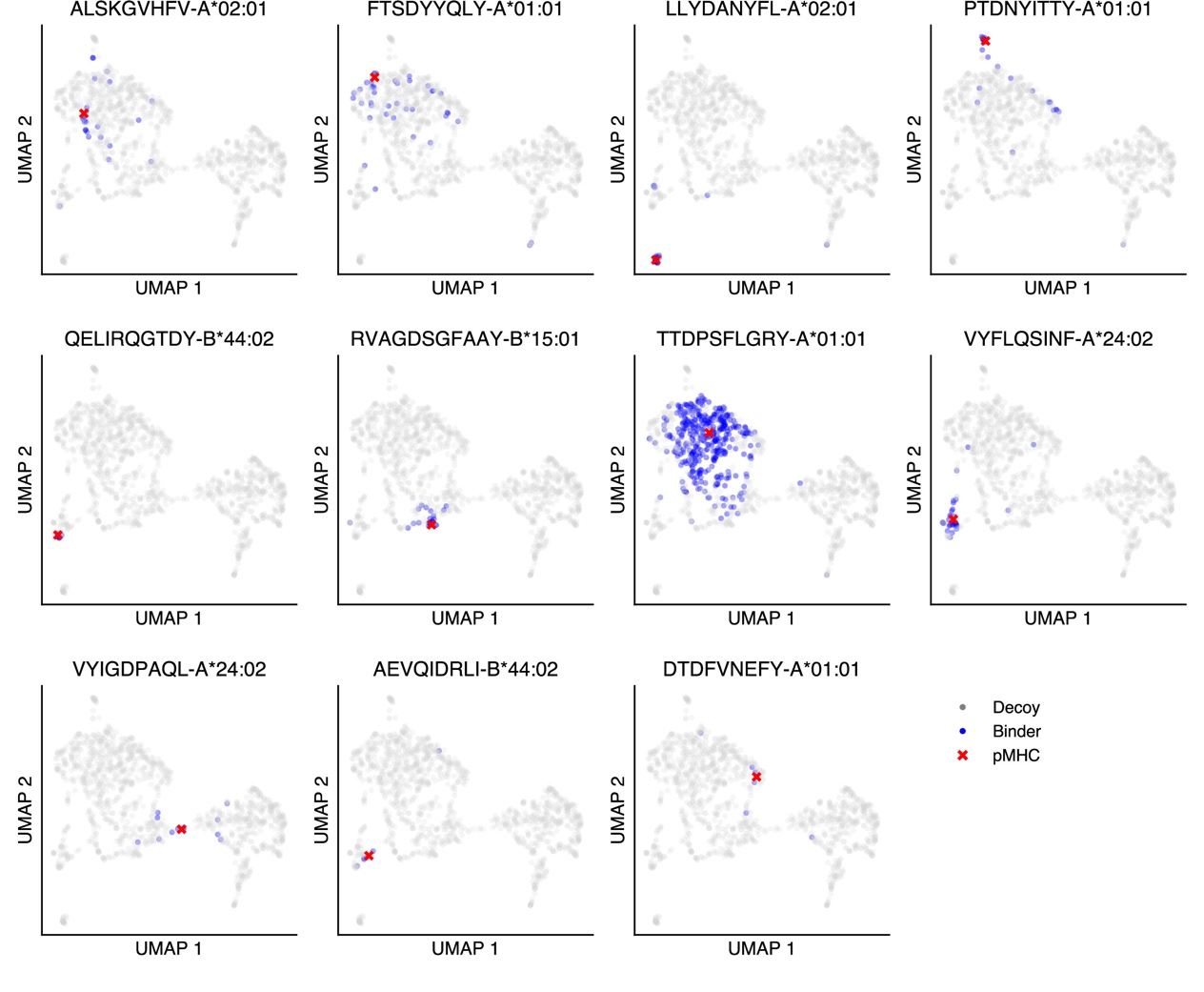

**Supplementary Fig. 5** Co-embedding visualization of SARS-CoV-2 antigen(non-spike)-specific TCRs.

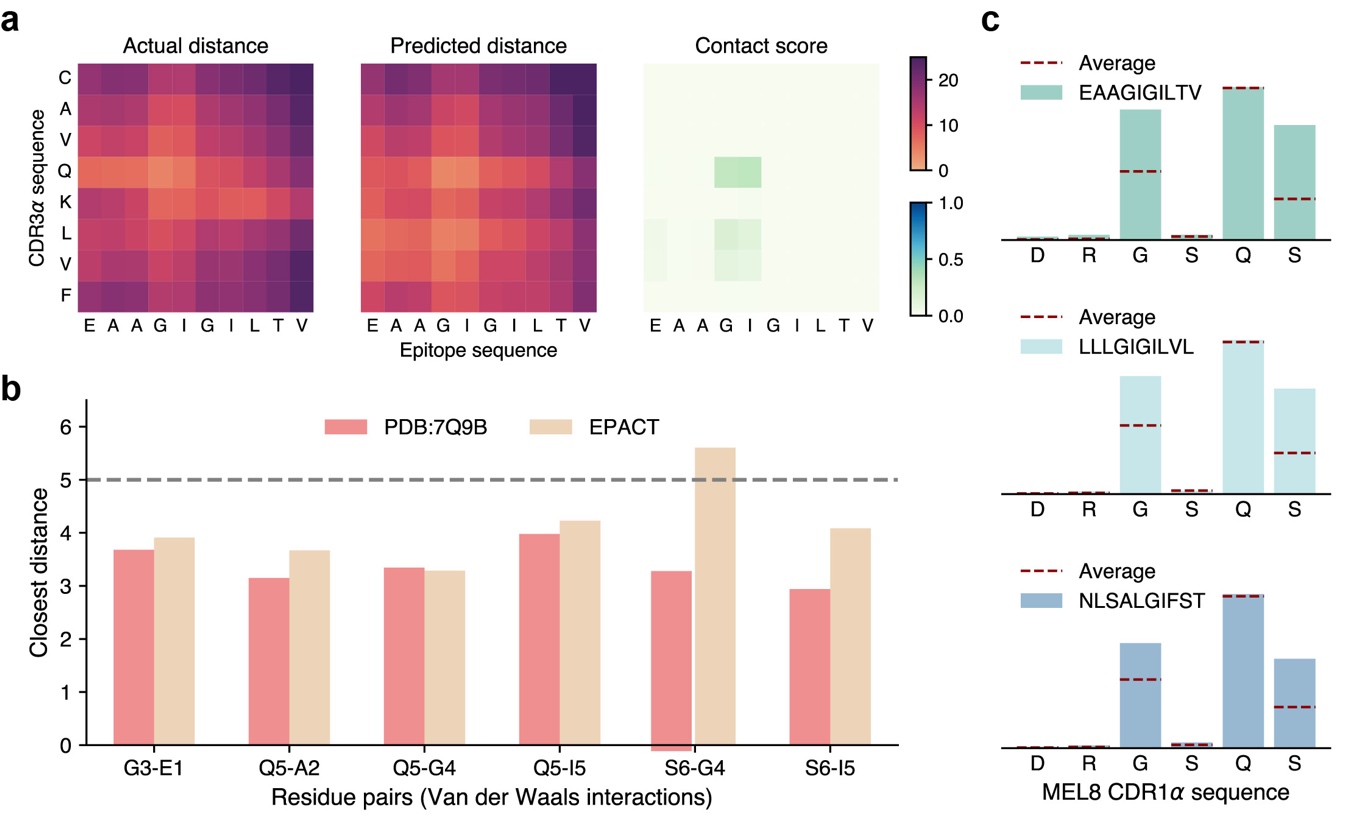

**Supplementary Fig. 6** EPACT recognizes the interacting residues between the MEL8 CDR1α sequence and cognate epitopes.

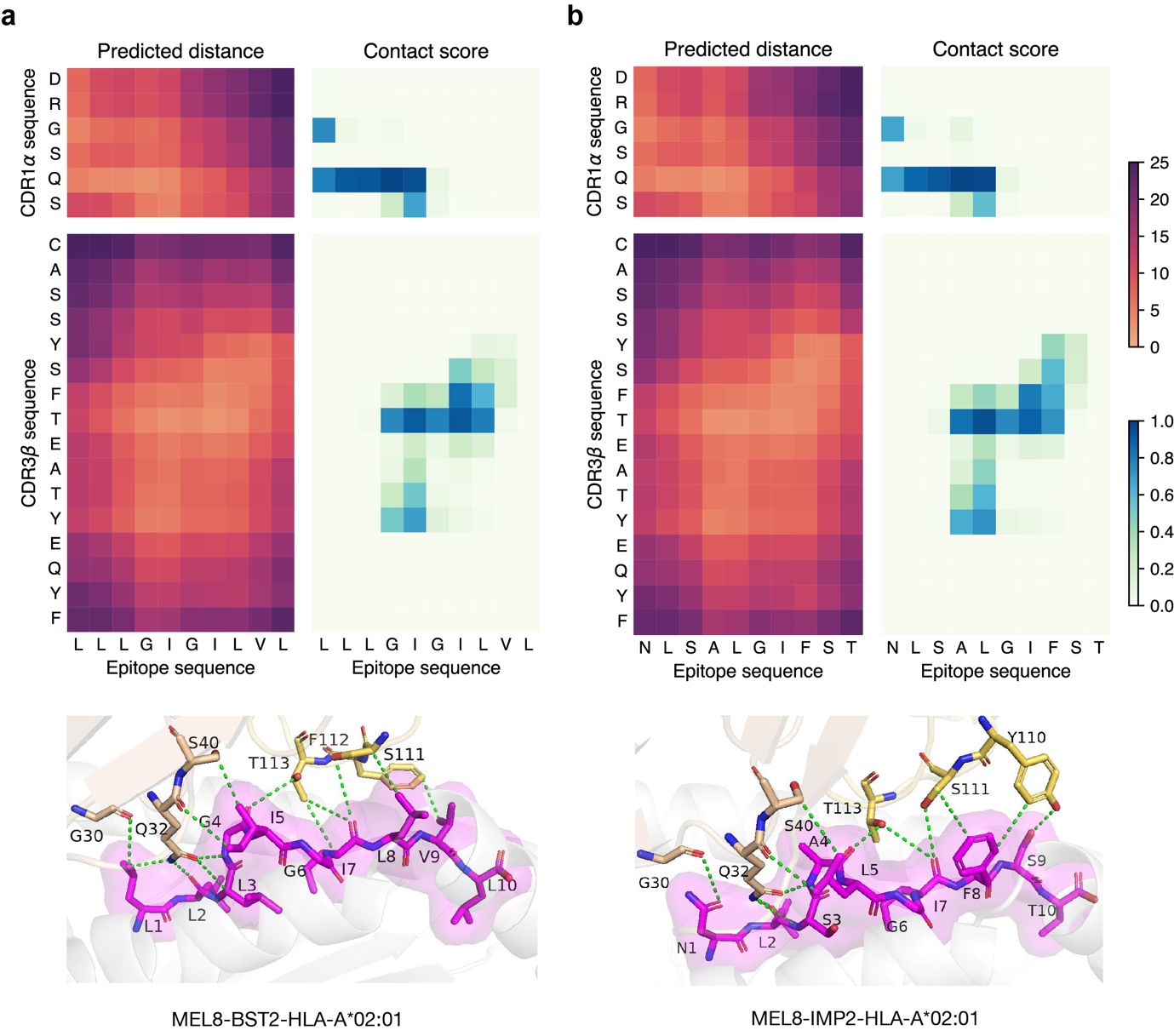

**Supplementary Fig. 7** Mutual corroboration of EPACT-predicted interactions and structure modeling of MEL8 TCR-pMHC complex.

Supplementary methods

Dataset pre-processing

Pre-processing of TCRαβ-pMHC recognition dataset

We restricted the length of peptides to 8-12 amino acids, which is within the typical length of CD8^+^ T-cell epitopes, and removed the peptides containing non-IUPAC amino acids. Many CDR3β sequences in the IEDB dataset lacked the starting cysteine (C). To adjust the sequence distributions of different data sources, we added the starting C for the CDR3 sequences not starting with cysteine (C) and the ending F for those not ending with phenylalanine (F) or tryptophan (W). We also filtered the CDR3 sequences by the criteria of IUPAC standard amino acids and sequence length between 10-25 amino acids. We then kept the epitopes having over five binding TCRs in the combined dataset, resulting in 24,500 TCRαβ-pMHC binding pairs spanning 250 CD8^+^ T-cell epitopes.

However, CDR3αβ sequences paired with the KLG peptide (KLGGALQAK) in the 10X dataset made up 48.7% of the TCRαβ-pMHC binding data. To address the imbalance of TCR binding specificity, we employed MMseq2^1^ cluster to retain representative TCR(CDR3)αβ sequences under different similarity thresholds. For CDR3αβ sequences paired with the KLG peptide, the minimum sequence similarity was set to 0.6, while a threshold of 0.9 was used for CDR3αβ paired with other epitopes. Finally, we built a test dataset comprising CDR3αβ-pMHC binding pairs that only occurred in the VDJdb dataset (1,147 positive pairs of 55 epitopes), and the remaining data (11,053 positive pairs of 240 epitopes) was used for model training.

Annotation of CDR1 and CDR2 sequences

Over 90% of the TCRαβ-pMHC binding pairs have annotated V and J genes. We downloaded the reference amino acid sequences of human TRAV, TRAJ, TRBV, and TRBJ alleles from the IMGT/GENE-DB84 database. Since the TRAV and TRBV alleles were presented in the form of “F+ORF+in-frame P with IMGT Gaps”, we extracted the CDR1 and CDR2 sequences of TCR alpha and beta chains according to the position of the IMGT-annotated sequences (CDR1:27-38, CDR2:56-65). Finally, we added CDR1 and CDR2 sequences to the TCRs in our pre-trained corpus and TCR-pMHC binding specificity data. Thus, we built a TCRαβ-pMHC recognition dataset containing all six CDR loops of TCR (9,936 positive pairs of 179 epitopes for training and 1,176 positive pairs of 62 epitopes for testing) in a similar way that mentioned in the previous section.

Generation of negative TCRαβ-pMHC pairs

To generate the negative TCRαβ-pMHC pairs for training and evaluation, we decided to sample five times “non-binding” TCRs with no replacement from those not paired with that peptide in the training or test dataset for each pMHC complex. Neither CDR3α nor CDR3β sequence of the “non-binding” TCRs appeared in the corresponding binding TCRs. Hence, we obtained a training dataset of 9,936(11,053) positive and 49,133(55,265) negative pairs for predicting TCR(CDR3)αβ-pMHC recognition. The VDJdb test dataset contained 1,176(1,147) positive and 5,837(5,564) negative pairs. Non-binding TCRs for the SARS-CoV-2 epitopes were sampled from the TCR repertoires (123,831 unique TCR clonotypes) of healthy human samples in 10X Genomics Datasets. The ratio of binding and non-binding TCRs was set to 1:5.

Pre-trained model development

Peptide language model

A Transformer-based protein language model^2^ was employed to capture the underlying representation of human peptides. The peptide sequences were randomly masked at a ratio of 0.15 and then input into an embedding layer to transform amino acid sequences into numeric embeddings. A learnable positional encoding layer was used to add position information to the sequence embeddings. After that, multiple transformer blocks comprising skip-connected multi-head attention layer and feed-forward network were adopted to learn the semantic relationships between amino acids. At last, a language modeling (LM) head consisting of a multi-layer perceptron (MLP) and softmax activation function was employed to predict the amino acid at the masked positions. The number of transformer layers, attention heads, and the hidden dimension size were set to 6, 4, and 512, respectively. A masked language modeling (MLM) loss was optimized using an AdamW optimizer^3^ with a learning rate of 1e-4 for 200 epochs. Besides the LM head, an additional MLP was employed to predict the Atchley Factors of the masked amino acids to help encode the biophysical and biochemical properties of amino acids. The mean square error (MSE) loss of predicted Atchley Factors was also involved in parameter optimization.

$$\mathcal{L}_{\mathrm{LM}}=\mathcal{L}_{\mathrm{MLM}}+ \mathcal{L}_{\mathrm{MSE}}=-\sum_{\hat{x}\in m\left( x \right)} \log p\left( \hat{x} | x_{\backslash m\left( x \right)} \right) + \varphi\left( \hat{y}-g\left( \hat{x} \right) \right)^{2},$$

where $m(x)$and $\backslash m(x)$ denote the masked and unmasked indices of the protein sequence, $\hat{y}$ denotes the predicted Atchley Factors, $g\left( \cdot\right)$ represents the mapping from amino acids to Atchley Factors, and $\varphi$ is the weighting factor of the MSE loss.

Paired TCR language model

A similar architecture as the peptide language model with some modifications was adopted to pre-train the representations of TCRαβ sequences. Firstly, paired CDR1α, CDR2α, CDR3α, CDR1β, CDR2β, and CDR3β sequences were concatenated in order (the CDR3 model only input the concatenated sequence of CDR3α and CDR3β). The CDR3 sequences were randomly masked at a ratio of 0.2, while other CDR segments were masked at a ratio of 0.1. Due to the variable lengths of CDR (especially CDR3) sequences from different TCRs, CDR segment-specific learnable positional encoding was used instead, defining the tokens’ position for each CDR loop, respectively. The hyperparameters of the transformer backbone and the loss function are the same as those of the peptide language model. An AdamW optimizer with a learning rate of 1e-4 was used to train the paired TCR(CDR3) language model for 100 epochs.

Peptide-MHC binding/presentation model

The amino acid sequences of the alpha chain of MHC class I molecules were padded to 366 amino acids and then converted into one-hot encoding(20d), BLOSUM50^4^ encoding(20d), and Atchley Factors(5d) to depict the biological features of MHC molecules. MHC feature embeddings(45d) were transformed to 256d vectors by 1D convolution and then input into an encoder network of six residual 1D convolutional blocks. To elucidate the residue-level interactions between peptides and MHC molecules, a dual cross-attention (DCA) module was utilized to integrate the residue-level information from pre-trained peptide representations into MHC embeddings and vice versa. DCA consisted of two symmetric multi-head cross-attention layers in which the query came from one sequence modality while the key and value were from the other. After that, crucial residues possibly contacting peptide residues of the MHC molecules were underscored through the attention pooling layer, and the pooled representations were concatenated with the classification [CLS] embeddings of the peptides. Finally, an MLP with a sigmoid activation function was employed to predict the binding or presentation of input peptide-MHC pairs. The MSE and Focal losses^5^ were used for binding affinity (BA) and eluted ligands (EL) data, respectively. In the training stage, input peptide sequences were masked for data augmentation, and an LM head was employed to recover the masked amino acids.

$$\mathcal{L}_{\mathrm{BA}}=\sum_{i} \left( f_{\mathrm{BA}}\left( \mathrm{peptide}_{i},\mathrm{MHC}_{i} \right)-t_{i} \right)^{2}+\lambda\mathcal{L}_{\mathrm{MLM}},$$

$$\mathcal{l}_{\mathrm{BCE},i}=-t_{i}\log f_{\mathrm{EL}}\left( \mathrm{peptide}_{i},\mathrm{MHC}_{i} \right)-\left( 1-t_{i} \right)\log\left( 1-f_{\mathrm{EL}}\left( \mathrm{peptide}_{i},\mathrm{MHC}_{i} \right) \right),$$

$$\mathcal{L}_{\mathrm{EL}}=\sum_{i} {\alpha\left( t_{i} \right)\left( 1-\exp\left( -\mathcal{l}_{\mathrm{BCE},i} \right) \right)^{\gamma}\mathcal{l}}_{\mathrm{BCE},i}+\lambda\mathcal{L}_{\mathrm{MLM}},$$

where $f_{\mathrm{BA}}$ and $f_{\mathrm{EL}}$ denote the output of pMHC binding and presentation model, respectively. $t_{i}$ denotes the prediction target, and $\alpha$, $\gamma$ are hyperparameters of Focal loss. Parameters of the pre-trained peptide language model were fixed during training, and an AdamW optimizer with a learning rate of 5e-4/1e-4 was used to train the pMHC binding/presentation prediction model. Unlike the training process, the complete sequences of peptides were input into the model during evaluation.

Benchmarking methods

Epitope presentation prediction

NetMHCpan-4.1^6^ is a pan-allelic model that simultaneously predicts peptide-MHC binding affinity and eluted ligands. It ensembles the predictions of 100 single-layer neural networks that receive a 9-mer binding core of peptide and a 34-mer MHC class I pseudo-sequence. The model outputs contain raw scores in the range of [0-1] and percentage ranks in the range of [0-100]. BigMHC^7^ introduces a new MHC pseudo-sequence representation derived from multiple sequence alignments. The model employs a wide LSTM architecture that increases the window size of one LSTM cell and anchor blocks to focus on anchor site binding residues. TransPHLA^8^ and MixMHCpred-2.1^9^ were also included in the performance comparison.

Binding specificity prediction

ERGO-II^10^ is based on the dual encoding of TCR and peptide. An LSTM network encodes peptide sequences while CDR3αβ sequences are encoded by autoencoder or LSTM. V and J genes of TCR and MHC alleles are considered categorical features. All feature encodings are input into an MLP classifier to predict binding specificity. NetTCR-2.0^11^ is a 1D CNN model that inputs the peptide, the CDR3α, and CDR3β region to indicate whether a given TCR can bind to a specific peptide. BLOSUM50 matrix is used for feature encoding, and the convolutional outputs are concatenated to produce the binding probability. NetTCR-2.2^12^ further combines the architectures of pan-specific and peptide-specific models and integrates CDR1 and CDR2 sequences into the CNN model. STAPLER^13^ adopts a shared TCR and peptide language model to learn the representations of TCR-peptide pairs exploiting a large pool of CDR3αβ sequences and 9-mer peptides. The pre-trained STAPLER model is fine-tuned on labeled pairs of CD8^+^ T cell-derived TCRαβ sequences and peptides that bind to MHC-I molecules. TULIP-TCR^14^ presents an unsupervised approach to model the relationship between CDR3α, CDR3β, epitope, and MHC via an encoder-decoder language model. It is only trained on interacting TCR-pMHC pairs and defines conditional probability for prediction. MixTCRpred^15^ utilizes transformer encoders to incorporate paired inputs of TCRαβ (CDR1, CDR2, and CDR3) sequences. It provides epitope-specific TCR-pMHC interaction models for 146 known viral and cancer epitopes presented by MHC class I and II alleles.

Interaction conformation prediction

TEIM-Res^16^ manipulates the interaction features from CDR3β and epitope sequences using an interaction extractor that employs 2D CNN modules to extract pairwise residue interaction information. The model is pre-trained on TCR binding specificity data and fine-tuned via a residue-level prediction module that outputs the predicted distance matrix and contact matrix. The average baseline for interaction prediction calculates the average distance and contact matrices of training samples as the prediction for any validating CDR3β-epitope pairs.

References

1.	Steinegger, M. & Söding, J. MMseqs2 enables sensitive protein sequence searching for the analysis of massive data sets. *Nat Biotechnol* **35**, 1026-1028 (2017).

2.	Lin, Z. et al. Evolutionary-scale prediction of atomic-level protein structure with a language model. *Science* **379**, 1123-1130 (2023).

3.	Loshchilov, I. & Hutter, F. Decoupled weight decay regularization. Preprint at <https://doi.org/10.48550/arXiv.41711.05101> (2017).

4.	Henikoff, S. & Henikoff, J.G. Amino acid substitution matrices from protein blocks. *Proc Natl Acad Sci U S A* **89**, 10915-10919 (1992).

5.	Lin, T.-Y., Goyal, P., Girshick, R., He, K. & Dollár, P. in Proceedings of the IEEE international conference on computer vision 2980-2988 (2017).

6.	Reynisson, B., Alvarez, B., Paul, S., Peters, B. & Nielsen, M. NetMHCpan-4.1 and NetMHCIIpan-4.0: improved predictions of MHC antigen presentation by concurrent motif deconvolution and integration of MS MHC eluted ligand data. *Nucleic Acids Res* **48**, W449-W454 (2020).

7.	Albert, B.A. et al. Deep neural networks predict class I major histocompatibility complex epitope presentation and transfer learn neoepitope immunogenicity. *Nat Mach Intell* **5**, 861-872 (2023).

8.	Chu, Y. et al. A transformer-based model to predict peptide–HLA class I binding and optimize mutated peptides for vaccine design. *Nat Mach Intell* **4**, 300-311 (2022).

9.	Gfeller, D. et al. Improved predictions of antigen presentation and TCR recognition with MixMHCpred2.2 and PRIME2.0 reveal potent SARS-CoV-2 CD8+ T-cell epitopes. *Cell Syst* **14**, 72-83.e75 (2023).

10.	Reiser, J.-B. et al. CDR3 loop flexibility contributes to the degeneracy of TCR recognition. *Nat Immunol* **4**, 241-247 (2003).

11.	Montemurro, A. et al. NetTCR-2.0 enables accurate prediction of TCR-peptide binding by using paired TCRα and β sequence data. *Commun Biol* **4**, 1060 (2021).

12.	Jensen, M.F. & Nielsen, M. NetTCR 2.2 - Improved TCR specificity predictions by combining pan- and peptide-specific training strategies, loss-scaling and integration of sequence similarity. *eLife* **12**, RP93934 (2023).

13.	Bjørn, P.Y.K. et al. STAPLER: Efficient learning of TCR-peptide specificity prediction from full-length TCR-peptide data. *bioRxiv*, 2023.04.25.538237 (2023).

14.	Barthelemy, M.-P., Christoph, F., Martin, W., Aleksandra, M.W. & Thierry, M. TULIP — a Transformer based Unsupervised Language model for Interacting Peptides and T-cell receptors that generalizes to unseen epitopes. *bioRxiv*, 2023.07.19.549669 (2024).

15.	Giancarlo, C. et al. Deep learning predictions of TCR-epitope interactions reveal epitope-specific chains in dual alpha T cells. *bioRxiv*, 2023.09.13.557561 (2023).

16.	Peng, X. et al. Characterizing the interaction conformation between T-cell receptors and epitopes with deep learning. *Nat Mach Intell* **5**, 395-407 (2023).
